# Bacterial assembly in the switchgrass rhizosphere is shaped by phylogeny, host genotype, and growing site

**DOI:** 10.1101/2023.11.15.567221

**Authors:** Jeremy Sutherland, Terrence H. Bell, Stacy Bonos, Christopher Tkach, Julie Hansen, Ryan Crawford, John E. Carlson, Jesse R. Lasky

## Abstract

- Since microbial traits are conserved at different taxonomic levels, plant hosts may influence microbiome composition differently at different levels to broadly promote or resist microbiota with traits that impact host fitness. We tested this hypothesis by assessing signals of host genetic influence on bacterial composition in the switchgrass rhizosphere using 128 genotypes in dissimilar growing sites.
- We employed three common gardens, combined with host genetic mapping, 16S rRNA gene sequence analysis, hierarchical modeling, tests of phylogenetic conservation of host influence, and genome-wide association analyses to determine the contributions of host genetics in shaping rhizosphere bacterial composition at different taxonomic levels.
- Modeling bacterial assembly showed that growing site was a strong factor shaping bacterial composition in the rhizosphere, though host genetic influence played a significant role. The heritability of bacterial abundance was strongest at the genus level. Phylogenetic signal for heritability was detected within the bacterial phylogeny but conserved clades differed between common gardens. We identified shared host genetic variants associated with bacterial abundance and host traits related to plant metabolism.
- Our results suggest further investigation is required regarding the genotype-by-environment-by-microbiome relationship to elucidate the factors shaping rhizosphere microbiome composition and the agroecological dynamics shaping plant phenotype.

## Introduction

Understanding how host-associated microbiomes assemble is an important issue for basic and applied community ecology, population biology, and plant breeding. From a community perspective, microbiome assembly presents a challenge, in part because microbial diversity is often even more diverse than paragons of community ecology, such as tropical tree communities (Prosser & Martiny, 2020). A wide range of abiotic constraints and biotic interactions are known to drive microbiome composition, but the largely unknown role of host physiology, evolution, and ecology in driving microbiome composition begs for more inquiry at the intersection of population and host organismal biology (Brunel et al., 2020). Furthermore, the importance of host-microbe interactions influencing host performance and ecosystem function suggests a high potential payoff for leveraging knowledge for applied purposes, such as plant domestication (Pérez-Jaramillo et al., 2016) and agricultural management strategies (Mahmud et al., 2021) that reduce the need for costly soil inputs.

Environmental effects, ecological drift, and dispersal limitations can influence the assembly of host-associated microbiomes at different spatial and temporal scales. For example, abiotic features of the environment can directly shape host-associated microbiome composition and can be a strong predictor of microbiome composition at macroscales, especially for microbiota found outside the plant (e.g., the rhizosphere) (Brunel et al., 2020). As a result, the presence and abundance of microbial taxa that associate with a given host can vary substantially across the host’s range (Serna-Chavez et al., 2013), meaning that different microbial strains, or coarser taxonomic groups or microbes, may be responsible for conserved interactions with hosts at different locations (Compant et al., 2010; Mansfield et al., 2012). This variability in host-microbe interactions among locations complicates our ability to consistently detect signals of host genetic influence on specific taxa through space (Fierer et al., 2013; Peiffer et al., 2013; Ruhl et al., 2022). To overcome this limitation, we employed three common garden experiments and genetic mapping to disentangle host genetic impacts on microbiome composition from site interactions.

Moreover, in community ecology, substantial progress has been made by considering community assembly in light of the phylogenetic relationships among species (Kembel & Hubbell, 2006). This approach is based on the hypothesis that traits determining species’ niches are phylogenetically conserved (Ackerly, 2003). Therefore, we hypothesized that if the microbial response to host traits and the abiotic environment is phylogenetically conserved (Martiny et al., 2015), then host genetic influences on microbiomes may respond to and affect microbiomes within clades where host fitness-related traits are more common while remaining relatively impartial to strain-level differences observed between sites. For example, if the consumption of a specific organic compound exuded by plants was preferred among members of a particular genus of soil bacteria (Zhalnina et al., 2018), then multiple strains within that genus might consume these compounds across different locations. This relationship may then directionally impact the next generation of hosts, resulting in a selective feedback loop over time (Hu et al., 2018). To assess this, we examined host-bacterial relationships across 128 clonal switchgrass genotypes with diverse life histories at three common gardens across the Northeastern United States, specifically assessing host genomic influence on root-associated microbiome composition at multiple levels of bacterial taxonomy.

Switchgrass (*Panicum virgatum*) life history defines most of the phenotypic differences between ecotypes (upland and lowland), cytotypes (tetraploid, octoploid, hybrid), and related genotype groups (North, South, East, West, and Northeast (N, S, E, W, NE)) (Lowry et al., 2015). Notably, these categories are linked to differences in root architecture and exudation (Stewart et al., 2017), which can influence microbial composition in the rhizosphere (Ulbrich et al., 2021). Our previous study demonstrated host genetic influence on switchgrass rhizosphere bacterial composition at a single site and related those influences to switchgrass life history traits (Sutherland et al., 2022). Yet much remains unknown regarding the genetic and phenotypic relationships between switchgrass hosts and their root-associated microbiomes across different growing sites and microbial taxonomy (Hestrin et al., 2021). Here, we examined two host traits that are of great interest to advancing the breeding and production of switchgrass in the northeastern United States: biomass yield and resistance to anthracnose disease (*Colletotrichum cereale)*, a common fungal pathogen in switchgrass. The interaction of host genetic variation, the environment, and microbiome composition influences on performance traits such as yield and disease resistance could be applied to breeding efforts that aim to influence microbiome composition and improve desirable host traits simultaneously. Here, we examined the host genotype-by-environment-by-microbiome interactions influencing these traits (anthracnose symptom severity and biomass yield) between three distinct environments in the Northeastern U.S. using a large panel of host switchgrass genotypes.

Toward this goal, we asked the following questions:

1. How does the growing site impact host genetic influence on microbiome composition in the switchgrass rhizosphere? The interaction between site-specific environmental factors (e.g., resource availability) and hosts could impact the host’s relative investment in microbiome composition. As a result, we might expect plant-associated microbiome diversity and composition to reflect a host’s relative resource needs in a given site. For example, soil conditions that promote the fast cycling of nutrients and high productivity can result in increased microbial diversity (Delgado-Baquerizo et al., 2017).
2. Is host genetic influence differentiated between bacterial taxonomic levels? If microbial traits are conserved phylogenetically, and those traits impact a host’s fitness in a given site, then host genetic influences on microbiomes may respond to and affect taxa differently throughout the bacterial phylogeny.
3. How do the interactions between switchgrass, their associated microbiomes, and the abiotic environment impact host phenotype? If plant hosts influence their microbiomes by collectively responding to site-specific environmental conditions and associated microbiota (i.e., the genotype-by-environment-by-microbiome interaction), then the relative abundance of certain microbes under host genetic influence should correlate with beneficial host traits in the context of different site conditions.
4. Which regions of the switchgrass genome are simultaneously associated with bacterial abundances and host traits correlated with those abundances? Since multiple genes can influence microbiome composition (e.g., root exudates are derived from multiple biochemical pathways), variation both genome-wide and at specific loci should relate to the variability in microbiome composition, and we might also expect those genes to be involved in biochemical processes and cellular components that interface with microbiota in the rhizosphere. In other words, we might expect shared genetic variants associated with both host traits and with the relative abundance of bacteria strongly correlated with those traits.

## Materials and Methods

### Plant materials and experimental design

In 2016, switchgrass accessions (n = 525) were vegetatively propagated by collecting sections from basal clumps from a common garden provenance trial (Lu et al., 2013) in Ithaca, NY, USA, and transplanting to growing sites in 1) Freehold, NJ, USA, 2) Philipsburg, PA, USA and 3) a separate site in Ithaca, NY, USA. At each location, the plant root and tiller sections were again divided into three equal sections, and each was planted into one of three randomized plots, resulting in three replicates per site. Soil conditions differed substantially between sites, with sandy-loam, clay-loam, and rock/shale soil conditions at the NJ, NY, and PA sites, respectively. The NY and NJ sites were planted in May 2016, while plugs for the PA site were kept in a greenhouse for one year before planting to avoid die-offs due to the harsher soil conditions at the PA site, which was previously a strip mine. The PA site is representative of marginal agricultural land (Csikós & Tóth, 2023), which may be important for switchgrass production to avoid land competition between biomass and food crops (Hartman, Nippert, Orozco, & Springer, 2011).

### Switchgrass genotyping

We downloaded the raw exome capture reads from the National Center for Biotechnology Information under BioProject PRJNA280418 (Evans et al., 2018) for each genotype in the original provenance collection, and performed new read mapping and variant calling using version v5.1 of the switchgrass genome. Raw reads were trimmed and processed for quality using cutadapt v3.4 (Martin, 2011), and read quality was assessed before and after trimming using FastQC v0.11.9 (Andrews, 2020). Trimmed reads for each genotype were then aligned to the switchgrass reference genome v5.1 (https://phytozome-next.jgi.doe.gov/) using BWA-MEM v0.7.17-r1188 (Li, 2013) and piped to SAMTools v1.14 (Danecek et al., 2021) for conversion to BAM format and sorting. Read groups were assigned to BAM files from the same sample using SAMTools v1.14 – *addreplacerg* command. Binary variant call format (bcf) files were generated using the BCFtools v1.14 – mpileup command (Danecek et al., 2021). Allele counts were extracted for 147,180,132 single-nucleotide polymorphisms (SNPs). These markers were then filtered so that those remaining had minor allele frequency >= 0.01 and a mean read depth per sample between 15 and 600 calculated across all populations to avoid SNP markers that might be inaccurately mapped to repetitive portions of the genome.

### Rhizosphere sampling of switchgrass accessions

We selected 128 out of 384 switchgrass genotypes for rhizosphere soil sampling, aiming to maximize the genetic distance between genotypes among those with good post-transplant survival. Rhizosphere soil samples were collected over 13 days from July 12-24, 2019. Rhizosphere sampling was achieved using a 91.44 cm stainless steel soil sampler with a 2.54 cm outer diameter. Depending on the size of the plant, two to three cores were collected from within 10 cm of the base of the plant at a depth of ∼15 cm. Soil and fine roots were deposited into a polythene bag between each collection. Labeled bags were stored on ice during collection and transferred to −20**°**C storage at the end of each day.

Soil sampling resulted in 236 rhizosphere samples from NJ, 366 from NY, and 318 from PA (920 samples in total). Severe weather conditions within a narrow collection time window limited sampling efforts at the NJ common garden to two of three replicate blocks. Plant die-off was higher at the PA site compared to NY due to seasonal flooding in the third replicate block and harsher soil conditions.

### Soil DNA extractions and amplicon sequencing

To determine bacterial composition in the switchgrass rhizosphere during sampling, total DNA was extracted from ∼300 mg of soil using the NucleoSpin Soil 96 kit (Macherey-Nagel, Düren, Germany). Lysis was performed using the Buffer SL with Enhancer SX and on the FastPrep 24 homogenizer (MP Biomedicals, Santa Ana, CA, USA) at 4.0 m/s for 30 s. Briefly, amplicons targeting the V3-V4 region of the 16S rRNA gene were generated from all samples using universal bacterial primers 515F (5′-GTGYCAGCMGCCGCGGTAA-3′) and 806R (5′-GGACTACNVGGGTWTCTAAT-3′) (Apprill, Mcnally, Parsons, & Weber, 2015; Parada, Needham, & Fuhrman, 2016) with overhangs for attaching barcodes and standard Illumina overhang adaptors in a second PCR step (full protocol provided in the supplemental methods of Trexler & Bell (2019)). Sequencing was performed on an Illumina MiSeq using the 2 × 250 cycle v2 kit.

### Amplicon sequence analysis pipeline

Raw 16S rRNA gene sequences were analyzed using an adapted version of the *dada2* pipeline (Callahan et al., 2016) to generate amplicon sequence variants (ASVs). Reads were truncated above 240 bp and below 160 bp—sequences with any missing reads after truncation were discarded. Reads were then truncated at the first instance of a quality score less than 2. After truncation, reads that matched against the phiX Genome were discarded. Reads with higher than two expected errors were also discarded. Expected errors are calculated from the nominal definition of the quality score: EE = sum(10^(-Q/10)^). Filtered sequences were used to determine the error rate using the *dada2* function: *learnErrors().* The filtered sequences were then processed with the core sample inference algorithm, *dada()*, incorporating the learned error rates. Forward and reverse sequence reads were then merged. A sequence table was made, and chimeras were removed using the “consensus” method. Taxonomic assignments for the ASVs were defined against the SILVA 138 ribosomal RNA gene database (Quast et al., n.d.) using DECIPHER v2.14.0 (Wright, n.d.). The sequence variant and taxonomy tables were exported for downstream processing in *phyloseq* (McMurdie & Holmes, 2013).

In *phyloseq*, ASVs designated “NA” at the phylum level, non-Bacterial entries at the domain level, and those associated with chloroplasts and mitochondria were removed from the taxonomy file before further processing. Finally, we also removed samples with no ASVs remaining after pruning. Rarefying samples has effectively reduced false discovery rates when large differences exist between the average sample library size (Schloss & Arbor, 2023; Weiss et al., 2017). Our data were characteristic of this scenario. Therefore, we rarefied samples to 500 sequences to retain as many samples as possible while maintaining a sufficient sampling depth to detect the most prevalent taxa comprising the switchgrass rhizosphere microbiome.

### Bacterial diversity and composition analyses

We generated alpha diversity metrics using the rarefied ASV dataset in *phyloseq* (McMurdie & Holmes, 2013). Statistical significance was calculated using the *vegan* (v2.5.6) package (Oksanen et al., 2019) and other core functions in R version 3.6.3 (Ihaka & Gentleman, 1996). The Shannon diversity index was calculated using the *estimate_richness()* function in *phyloseq* and then used to test for differences in alpha diversity between genotypes differing in ecotype, ploidy level, and genetic cluster groups using an analysis of variance (ANOVA) (R version 3.6.3).

We calculated an NPMANOVA using the *adonis2* function in *vegan* to test for significant differences between ecotypes, cytotypes, and genetic clusters on the overall rhizosphere microbiome composition using pairwise Bray-Curtis dissimilarities at the ASV level within and between common gardens. A principal coordinate analysis (PCoA) was performed in *phyloseq* to observe compositional differences between common gardens.

### Heritability of rhizosphere bacteria

Using the rarefied ASV dataset, we estimated broad-sense heritability by fitting normalized count data for taxonomic groups at each taxonomic level (i.e., phylum, class, order, family, genus) to factorial host genotypes at each site (i.e., PA, NJ, NY, ALL -all genotypes in study). We used the *lm()* function in the *stats* package (v4.2.1) in R to estimate broad-sense heritability. Weather conditions and time constraints impaired rhizosphere sampling at the NJ common garden, resulting in missing data points for most of the third replicated plot. We removed the third replicate for each genotype at all sites to better reflect the sampling effort at the NJ common garden.

To test for host genetic influence on microbe traits, we treated heritability as a bacterial ‘trait’ and determined the phylogenetic signal based on spatial autocorrelation between heritability estimates and their respective positions within the bacterial phylogenetic tree (i.e., Local Moran’s *I*). We obtained phylogenetic trees from the Web of Life: Reference Phylogeny for Microbes, release 1 (Apr 5, 2019), built using 10,575 genomes and 381 genes (Zhu et al., 2019) and determined phylogenetic signal for heritability using the *phylosignal* package (v1.3) in R (Keck, Rimet, Bouchez, & Franc, 2016).

### Generalized linear mixed modeling and variance partitioning

We took a hierarchical species distribution modeling approach (here, across hosts) to identify common drivers of microbiome composition among diverse communities (Lasky et al., 2017). Here, we used generalized linear mixed effects regression (GLMER) to model rarefied bacterial ASV counts as a function of several potential drivers (covariates), where individual ASVs had random intercepts and covariate slopes. The covariates included growing site and the first two principal components of the switchgrass SNP PCA as random effects. We ran six models with ASV counts nested based on their respective taxonomy (i.e., phylum, class, order, family, genus, unnested). Switchgrass PCs were determined in R using a principal component analysis (PCA) of 196,772 switchgrass SNPs using the *rda*() function in *vegan* v2.6-2 (Oksanen et al., 2019). The top PCs of the switchgrass SNP data map well to genetic distances between genotypes and coarsely capture the allelic variation between switchgrass ecotypes (Supplemental Figure 1). Model estimates were obtained using GLMER with a log link function to model the Poisson distributed responses. Models were fit using the *glmm*() function in the *lme4* package (v1.1-30) in R.

We modeled multiple taxonomic levels to determine the relative host genetic effects on bacterial assembly between them. For each model nested among taxonomic levels, a full model using default *glmm*() settings was constructed with all the possible interactions among the main effects for growing site (NJ, NY, PA), and the first two principal components of the switchgrass SNP PCA. Additionally, two other models were constructed: the first included only growing site (Site-only) as a main effect, and the second included only the first two principal components of the switchgrass SNP PCA as the main effect (PC-only). To compare models downstream, we removed ASVs with an unknown classification at the genus level before modeling, ensuring all models shared the same number of observations. A likelihood ratio test of nested models (Hothorn, 2002) was employed to compare the hierarchically nested models (Full, Site-only, PC-only) to determine whether the additional complexity in the Full model was significantly more accurate than the reduced models. The Akaike information criterion (AIC) (Burnham & Anderson, 2002) was employed to identify the best model fit (Myung, Forster, & Browne, 2000).

### Host phenotyping and associations with microbiome abundances

Plants were harvested and weighed at each common garden in September-October 2021 with help and planning input from collaborators at Cornell University and Rutgers University, respectively. Plants were then transported to a facility at Cornell University in Ithaca, NY, USA for drying. Dried plant matter was then weighed and used to determine biomass yield. Before harvesting, symptom severity for anthracnose was recorded with “1” representing plants with virtually no disease symptoms, “2” representing plants with few characteristic symptoms (i.e., lesions), “3” showing moderate symptoms, “4” showing moderate-to-severe symptoms, and “5” showing severe symptoms, stunted growth and die off (<u>Appendix</u> Figure 1). Plant height, plant circumference, smut disease (*Tilletia maclaganii* (Berk.) Clint) symptom severity, and vigor were also collected during the 2021 growing season (<u>Appendix</u> Table 1). Pearson’s correlations, accounting for false discovery rate, between biomass yield, anthracnose disease severity, and microbial abundances were determined in R to identify potential bacteria that might relate to host traits.

**Figure 1:**
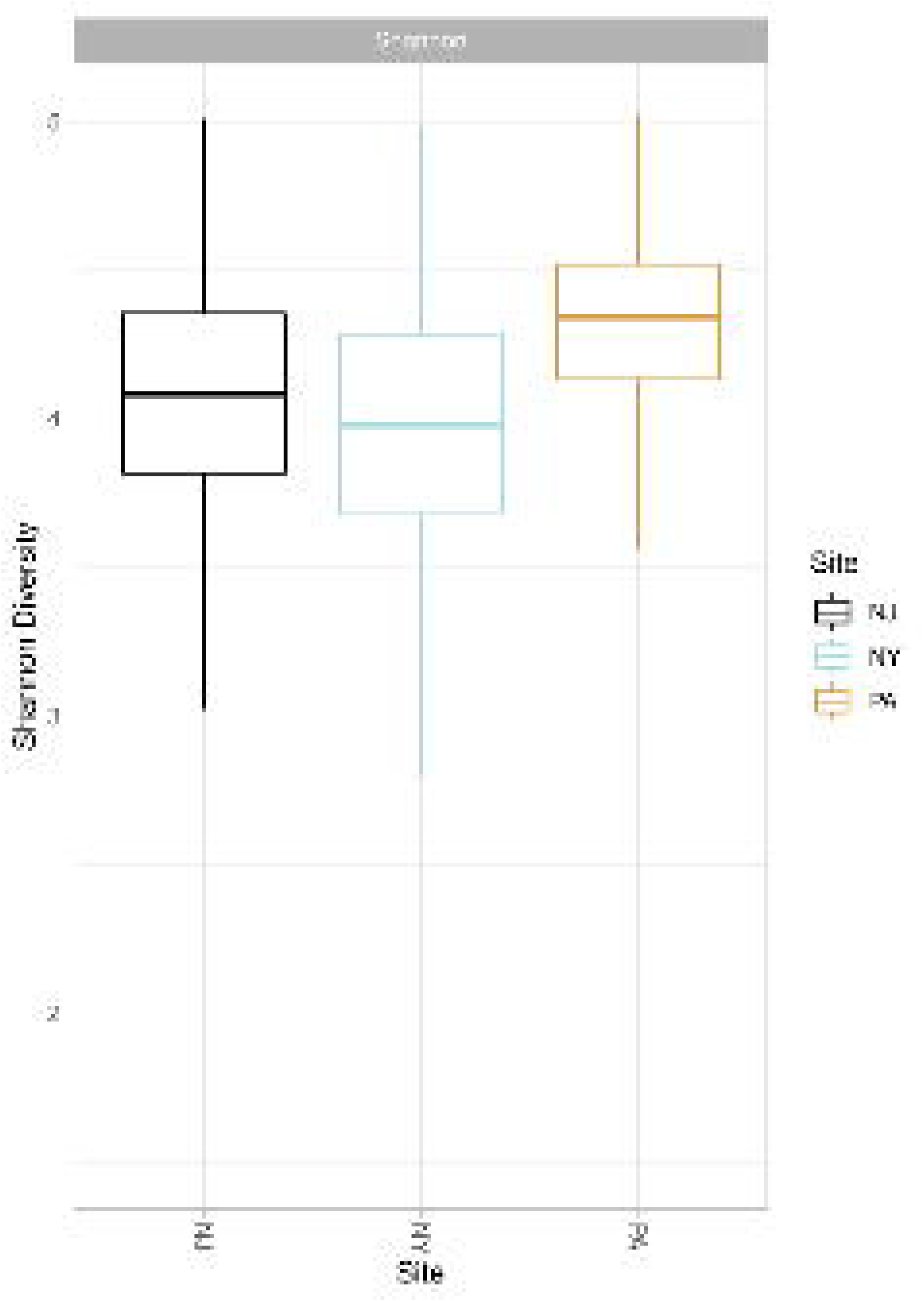
Bacterial Rhizosphere Alpha Diversity Between Sites. Shannon diversity index representing the change in bacterial diversity between common gardens in NJ (black), NY (blue), and PA (gold).

**Table 1:**
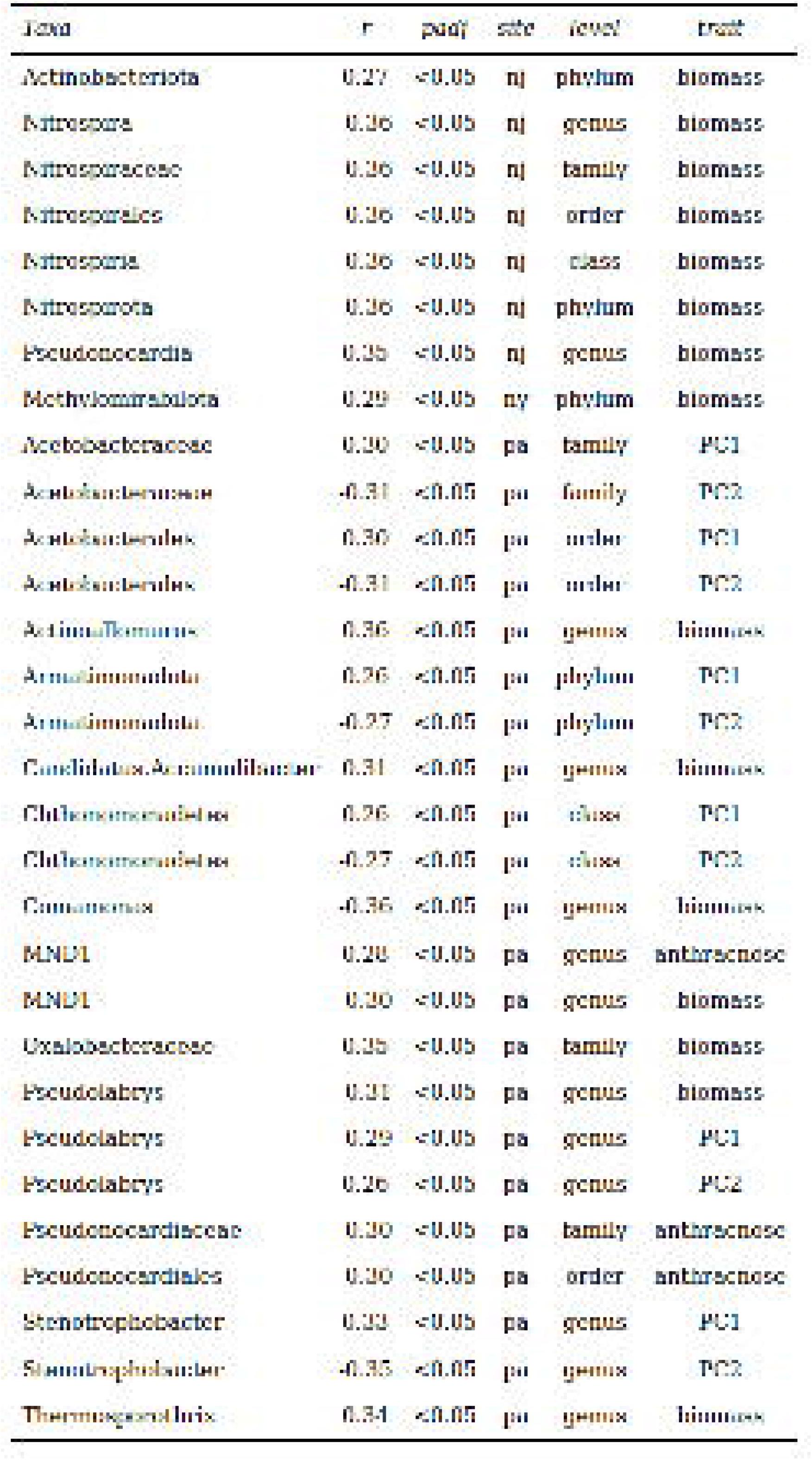
Significant Correlations Between Bacterial Abundances and Host Traits and SNP PCs. Table includes Pearson’s correlations (r) between the relative abundance of bacterial taxa at each site and host traits: biomass yield, anthracnose disease severity, and snpPCs 1 and 2 (trait), the significance cutoff, the adjusted significance cutoff, site, and taxonomic level.

### Genome-wide association study

To identify putative causal genes for host-associated variation in bacterial composition, we conducted several genome-wide association studies (GWAS). A GWAS was conducted using the *statgenGWAS* (Kang et al., 2010; Rossum, Kruijer, Eeuwijk, & Boer, 2020) package in R to identify specific loci in the switchgrass genome linked to two host phenotypic traits (biomass and anthracnose disease severity) and the abundance of bacterial taxa with significant associations to those traits at each common garden (NJ, NY, PA) (Table 1).

To determine any associated genes, a single trait GWAS was completed using a Generalized Least Squares (GLS) method for estimating the marker effects and corresponding p-values on scaled microbial taxa abundances. Our GWAS included random effects correlated according to the kinship matrix, calculated with the VanRaden method (VanRaden, 2008). We implemented a method to minimize False Discovery Rate (FDR) following the algorithm proposed by Brzyski et al. (Brzyski et al., 2017). In this method, SNPs were first restricted to those with a p-Value below 0.01. Then, clusters of SNPs were created using a two-step iterative process in which, first, the SNP with the lowest p-Value is selected as the cluster representative. This SNP and all SNPs correlating with this SNP at 0.9 or higher will form a cluster. A full description of the method can be found in Rossum et al. (2020).

We conducted a GWAS for the following taxa: *Actinobacteriota, Methylomirabilota, Nitrospiraceae, Oxalobacteraceae, Pseudolabrys, Pseudonocardia, Pseudonocardiaceae*. The relative abundance of these taxa was significantly correlated at one site or more with host traits: biomass yield and anthracnose disease severity. At times, the SNP effect sizes and p-values for taxa abundance were identical for multiple taxonomic levels. This can happen because a certain class, for instance, may contain only one family or genus. In these instances, we used the lowest level with a significant SNP association for the GWAS (e.g., *Nitrospiraceae* at NJ and *Pseduonocardiaceae* at PA). Additionally, although *Actinoallomurus, Candidatus Accumulibacter, Comamonas, Thermosporothrix,* and *Pseudonocardiales* were correlated with the two host traits at PA, they were disregarded due to few positive counts in the ASV dataset (i.e., zero-inflated) which likely resulted in spurious associations between their abundance and host traits.

## Results

For switchgrass genotyping, we identified 196,772 exome-captured SNPs (MAF >= 0.01) after filtering with *bcftools* (Danecek et al., 2021). For 920 initial rhizosphere soil samples across all three common gardens, we identified 18,539 unique amplicon sequence variants (ASVs) following initial quality filtering and sequence processing in *dada2* (Callahan et al., 2016). After sample pruning (removing non-bacteria from the taxonomy table) and rarefaction (500 sequences/sample) in *phyloseq*, 6,833 ASVs remained in 915 samples belonging to the bacterial domain, characterized by 23 phyla, 53 classes, 127 orders, 241 families, and 477 genera.

### Host effects on bacterial diversity and composition in the switchgrass rhizosphere

Using the Shannon Diversity Index to measure alpha diversity, we found no statistical differences in rhizosphere diversity between ecotypes, cytotypes, or genotype groups across sites. However, an analysis of variance (ANOVA) found that Shannon diversity was significantly different between common gardens (*p* = < 0.001). The PA garden, which is highly degraded with large amounts of rock and shale, exhibited the greatest rhizosphere alpha diversity on average. Samples from the NY site exhibited the least diversity on average (Figure 1).

We calculated an NPMANOVA to test for significant differences between overall rhizosphere microbiome composition within and between each common garden using pairwise Bray-Curtis dissimilarities of ASVs. We found significant differences in overall rhizosphere composition between common gardens (*R^2^* = 0.095, *p* = < 0.001, Figure 2) and between genetic clusters (N, S, E, W, NE) at the PA common garden (*R^2^* = 0.013, *p* = 0.012) but not at the other gardens. The non-parametric test revealed no significant differences between ecotypes (upland, lowland) or cytotypes (tetraploid, octoploid, hybrid) at any common garden.

**Figure 2:**
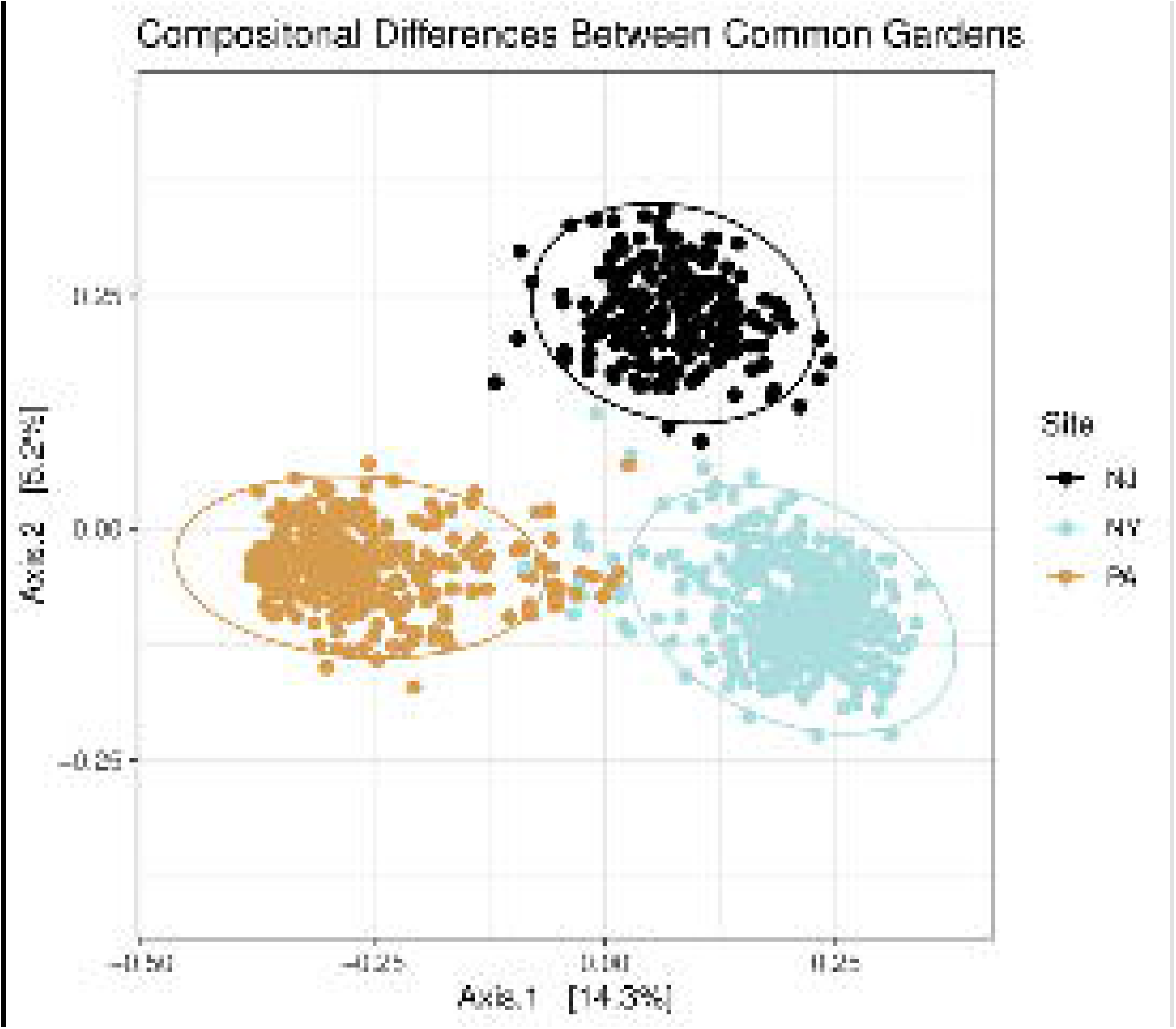
Principal Coordinate Analysis of Bacterial Composition between Sites. Principal coordinate analysis of Bray-Curtis dissimilarities in bacterial composition at the ASV-level, where points represent samples collected from common gardens in New Jersey (black), New York (blue), and Pennsylvania (gold). The corresponding ellipses represent a 95% confidence level for a multivariate t-distribution within each common garden.

### Heritability persisted across sites but differed between them

We estimated broad-sense heritability (*H^2^*) to describe the proportion of variance in relative microbial abundance due to switchgrass genetic effects at each common garden (NJ, NY, PA) and collectively for genotypes using samples collected across all sites (ALL). An analysis of variance found that mean differences in genus-level *H^2^* between the common gardens (NJ, NY, PA) were significantly different (*p* < 0.001) but not at other taxonomic levels. The highest *H^2^* estimates were at the NJ common garden (Figure 3). When we estimated *H^2^* for genus-level microbial abundances for genotypes across all sites (ALL), increasing the overall environmental variability, we saw a large reduction in the mean estimate. However, the non-zero mean estimate for ‘ALL’ indicates that host genetic influence transfers across these very distinct sites. As before, an analysis of variance found that mean differences in *H^2^* were significantly differentiated between all sites respectively (NJ, NY, PA, ALL) for bacterial genera present in the switchgrass rhizosphere (*p* < 0.001) (Figure 3).

**Figure 3:**
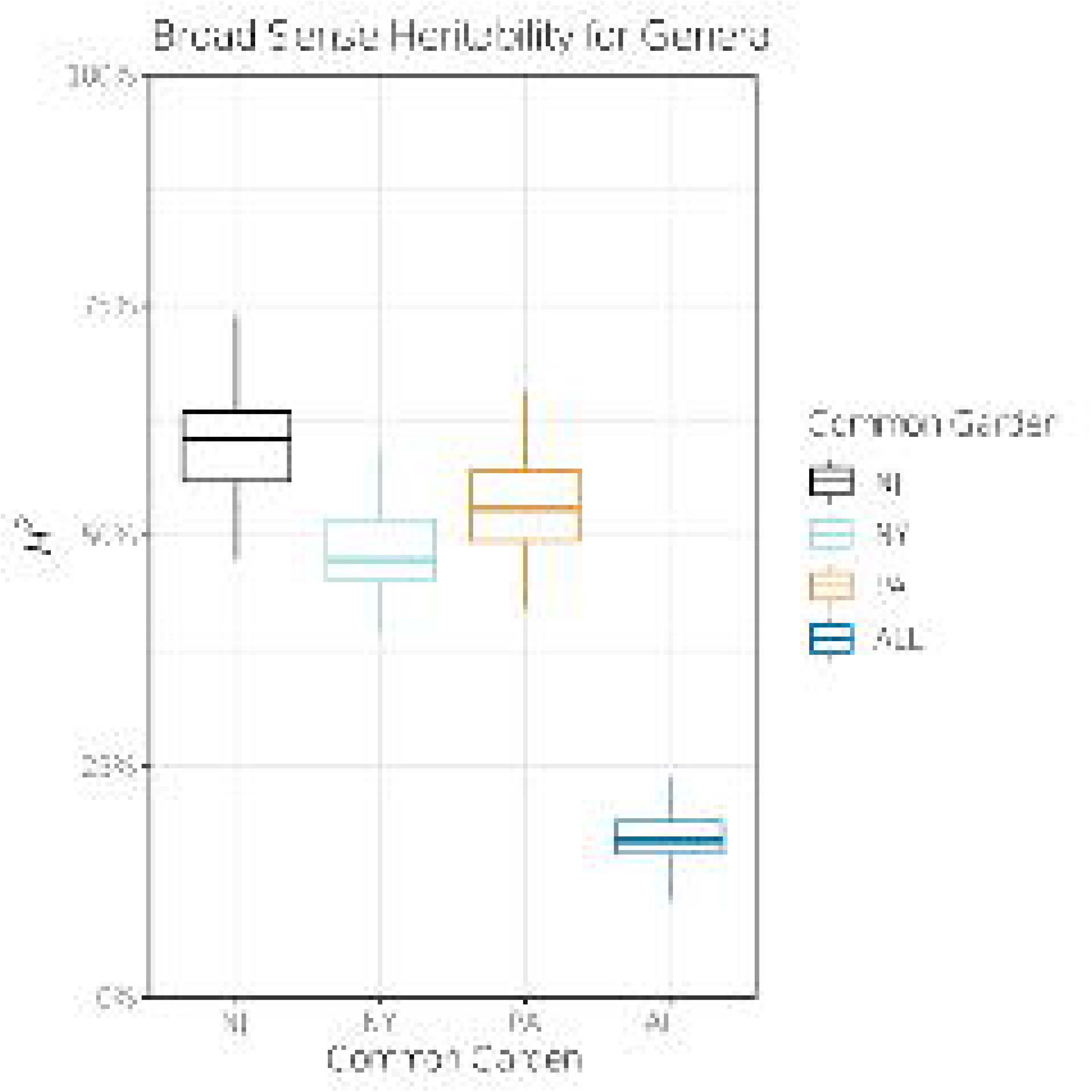
Broad Sense Heritability for Bacterial Genera. Broad-sense heritability estimates for genera at NJ (black), NY (blue), PA (gold), and ALL (dark blue).

### Bacterial community assembly is best explained by site and host genetics together

We implemented a hierarchical distribution modeling approach to test how the shared response to host genotype and growing site drove bacterial composition in the rhizosphere. We employed a generalized linear mixed model (GLMM) on the rarefied ASV counts to estimate the random effects of growing site (represented as a factor for each site: NJ, NY, PA) and the first two principal components of a switchgrass SNP-based principal component analysis. Likewise, variance components extracted from GLMMs effectively estimate the total variability accounted for by each specified source (i.e., host SNP PCs and growing site). A likelihood-ratio test was used to compare the goodness of fit of the statistical models at each taxonomic level. The Full model, compared to the Site-only or PC-only models, was more accurate than the reduced models in all cases (*p* < 0.001) (Table 3). Using AIC to estimate the relative quality of each Full model, we determined that unnested ASV counts were the best fit for the model, compared to nesting ASVs within their respective taxonomic rank (phylum, class, order, family, genus) (Table 3).

### Bacterial heritability was differentially conserved between sites

We estimated broad-sense heritability (*H^2^*) to describe the proportion of variance in bacterial abundance due to switchgrass genetic effects at different bacterial taxonomic levels. The highest *H^2^* estimates were generally for ASV counts grouped at lower taxonomic levels. For instance, the highest *H^2^* estimates for PA (0.796) and NJ (0.790) were at the genus level, while the highest estimate was at the family level for NY (0.647). For ASV counts grouped at the genus level, the mean *H^2^* estimates were 0.599 at NJ, 0.487 at NY, 0.534 at PA, and 0.169 across ALL (Figure 3).

Next, we tested for a bacterial phylogenetic pattern of heritability. Here, we used a phylogenetic correlogram for each taxonomic level to represent how heritability is autocorrelated at different phylogenetic distances. The Global Moran’s *I* Index was significant for heritability among genera at the NJ common garden (*p* < 0.05) and ALL (*p* < 0.01) but not at other taxonomic levels or common gardens. (Figure 4). This means that the distribution of heritability estimates with similar values was more clustered across the bacterial phylogeny of genera than expected at NJ and ALL but not for other sites or within other trees.

**Figure 4:**
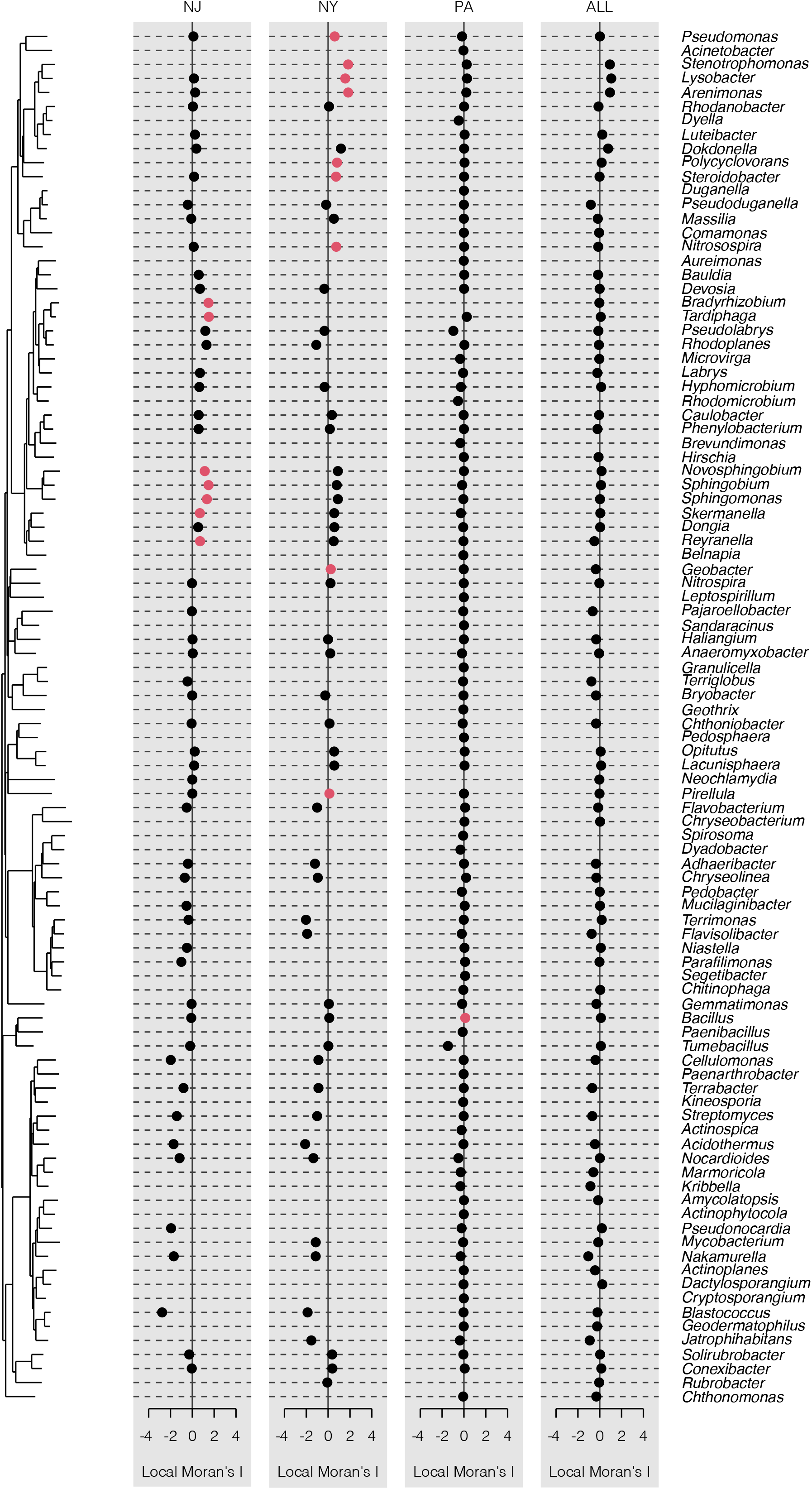
Local Indicator of Phylogenetic Association. Local Indicator of Phylogenetic Association (local Moran’s I) for genera autocorrelated with heritability in switchgrass. Significant associations between tree position and heritability are in Red.

We computed the local Moran’s *I* for each tip of each bacterial phylogenetic tree (phylum-genus) to determine the phylogenetic signal therein for each common garden. Here, a positive value for *I* indicates that a given taxon has neighboring taxa with similarly high or low heritability estimates (i.e., a cluster). A negative value for local *I* indicates that a taxon has neighboring taxa with dissimilar values (i.e., an outlier). In either instance, the p-value for the taxa must be small enough for the cluster or outlier to be considered statistically significant. We found that 17 genera formed clusters (NJ = 7, NY = 9, PA = 1, and ALL = 0) (Figure 4). There were no significant clusters or outliers at other taxonomic levels.

### Associations between bacterial abundances and host traits were significant

We determined associations between the relative abundance of bacteria (at different taxonomic levels) and four genetic or phenotypic characteristics of switchgrass: dried biomass weight (biomass), anthracnose disease severity (anthracnose), and the first two principal components of switchgrass SNPs (PC1, PC2). PC1 and PC2 relate to the genome-wide differences associated with upland and lowland ecotypes and genetic groups (Supplemental Figure 1). Mean biomass and anthracnose ratings were generally highest at NY (Supplemental Figure 3). Using Pearson’s correlations, we found 326 putative associations between these traits and the relative abundance of bacterial taxa among common gardens (NJ = 71, NY = 69, PA = 186). After adjusting for false discovery rate (FDR, 0.05), we confirmed 35 associations (NJ = 7, NY = 1, PA = 22, Table 1).

After FDR adjustment, all of the associations that were positively correlated with PC1 were negatively correlated with PC2. At NY, only one taxon positively correlated with biomass, compared to two at NJ and four at PA. At NJ, there were five taxa that negatively correlated with biomass. However, all five belonged to the phylum Nitrospirota. There were three taxa that negatively correlated with biomass at PA, and one of them, MND1, was also positively correlated with anthracnose. Finally, two taxa negatively correlated with anthracnose severity at PA. Both belonged to the order Pseudonocardiales. No taxa were associated with anthracnose severity at the NY and NJ sites, after FDR adjustment.

### A genome-wide association study revealed shared SNPs associated with host traits and bacterial abundances

We performed two genome-wide association studies. One for host traits (biomass and anthracnose) and one for bacterial abundances that correlated with those traits. Overlapping SNPs could reveal loci involved in traits that influence both. We found 646 SNPs significantly associated with the abundance of seven taxa (<u>See Materials and Methods</u>): NJ = 318, NY = 119, PA = 209. There were 610 SNPs significantly associated with Biomass Yield: NJ = 274, NY = 175, PA = 161; and 256 SNPs associated with Anthracnose Severity: NY = 85, NJ = 91, PA = 80. Altogether, fifteen SNPs belonging to thirteen genes were shared between the abundance of the seven taxa and host traits, one position was not annotated, and two positions were within the same gene (Table 2). An enrichment analysis showed no significant GO terms associated with these genes; however, most genes (n = 6) were associated with Gene Ontologies related to plant metabolism: GO:0008152: Pavir.1KG354700: *(1 of 6) K07877 -Ras-related protein Rab-2A (RAB2A),* Pavir.3KG080100: *K14484 -auxin-responsive protein IAA (IAA)*, Pavir.5KG119200: *PTHR21596//PTHR21596:SF26 - RIBONUCLEASE P SUBUNIT P38 // SUBFAMILY NOT NAMED*, Pavir.6KG071170: *K05279//K13066 - flavonol 3-O-methyltransferase (E2.1.1.76) // caffeic acid 3-O-methyltransferase (E2.1.1.68, COMT)*, Pavir.7KG162000: *PTHR24361//PTHR24361:SF337 - MITOGEN-ACTIVATED KINASE KINASE KINASE // SUBFAMILY NOT NAMED*, Pavir.7KG413400: *K10405 - kinesin family member C1 (KIFC1)*. A full list of genes and GO terms can be found in Table 2.

**Table 2:** GWAS Results: Significant SNPs associated with bacterial abundances correlated with host traits and host traits. Table includes significant SNPs associated with biomass yield and anthracnose disease severity, and the relative abundance of bacterial taxa in the switchgrass rhizosphere that correlate with those traits (p = <0.05, padj = <0.05), SNP name, chromosome, position, allele frequency, p value, effect size, effect size standard error, likelihood-ratio-based R2, LOD score, proportion of the variance explained by the SNP-computed as β2SNP∗var(SNP)/var(pheno, site, associated gene name and GO terms.

Overlapping SNPs: Bacterial Abundances and Host Traits
| trait | snp | chr | pos | allFreq | pValue | effect | effectSe | RLR2 | LOD | snpStatus | propSnpVar | site | gene | terms |
| --- | --- | --- | --- | --- | --- | --- | --- | --- | --- | --- | --- | --- | --- | --- |
| dry_biomass_g | Chr01K:38999450 | Chr01K | 38,999,450 | 0.1440 | < 0.001 | 0.8116 | 0.2387 | 0.0503 | 3.8570 | significant SNP | 0.0762 | PA | Pavir.1KG354700 | GO:0008152 |
| Pseudolabrys | Chr01K:38999450 | Chr01K | 38,999,450 | 0.1440 | < 0.001 | -0.8901 | 0.2514 | 0.0544 | 3.8969 | significant SNP | 0.0603 | PA | Pavir.1KG354700 | GO:0008152 |
| Pseudonocardia | Chr02N:6272636 | Chr02N | 6,272,636 | 0.3495 | < 0.001 | -0.7298 | 0.2190 | 0.0655 | 4.2864 | significant SNP | 0.0679 | NJ | Pavir.2NG061200 |  |
| dry_biomass_g | Chr02N:6272636 | Chr02N | 6,272,636 | 0.3520 | < 0.001 | -0.8940 | 0.2584 | 0.0520 | 3.9776 | significant SNP | 0.0640 | PA | Pavir.2NG061200 |  |
| dry_biomass_g | Chr03K:5792111 | Chr03K | 5,792,111 | 0.1410 | < 0.001 | -0.8142 | 0.2318 | 0.0590 | 4.2611 | significant SNP | 0.0826 | NY | Pavir.3KG080100 | GO:0043170, GO:0008152 |
| Pseudolabrys | Chr03K:5792111 | Chr03K | 5,792,111 | 0.1440 | < 0.001 | 1.3445 | 0.2903 | 0.0913 | 6.3619 | significant SNP | 0.1156 | PA | Pavir.3KG080100 | GO:0043170, GO:0008152 |
| dry_biomass_g | Chr04K:1218279 | Chr04K | 1,218,279 | 0.1880 | < 0.001 | -0.8534 | 0.2217 | 0.0640 | 4.8103 | significant SNP | 0.1014 | PA | Pavir.4KG003200 | GO:0006457, GO:0009987 |
| Pseudolabrys | Chr04K:1218279 | Chr04K | 1,218,279 | 0.1880 | < 0.001 | 1.1844 | 0.2489 | 0.0961 | 6.6959 | significant SNP | 0.1286 | PA | Pavir.4KG003200 | GO:0006457, GO:0009987 |
| anthracnose | Chr04K:37765796 | Chr04K | 37,765,796 | 0.3462 | < 0.001 | 0.2566 | 0.0714 | 0.0616 | 3.6336 | significant SNP | 0.1045 | NY | Pavir.4KG318200 |  |
| dry_biomass_g | Chr04K:37765796 | Chr04K | 37,765,796 | 0.3495 | < 0.001 | -0.9890 | 0.3221 | 0.0559 | 3.8302 | significant SNP | 0.0882 | NJ | Pavir.4KG318200 |  |
| anthracnose | Chr04K:47381132 | Chr04K | 47,381,132 | 0.1311 | < 0.001 | -0.3659 | 0.0971 | 0.0829 | 3.8053 | significant SNP | 0.0870 | NJ |  |  |
| dry_biomass_g | Chr04K:47381132 | Chr04K | 47,381,132 | 0.1311 | < 0.001 | 1.2428 | 0.3789 | 0.0635 | 4.3001 | significant SNP | 0.0666 | NJ |  |  |
| dry_biomass_g | Chr04N:34780024 | Chr04N | 34,780,024 | 0.4000 | < 0.001 | 0.7326 | 0.2205 | 0.0481 | 3.7038 | significant SNP | 0.1353 | PA | Pavir.4NG164500 |  |
| Pseudolabrys | Chr04N:34780024 | Chr04N | 34,780,024 | 0.4000 | < 0.001 | -0.8865 | 0.2308 | 0.0638 | 4.5097 | significant SNP | 0.1303 | PA | Pavir.4NG164500 |  |
| dry_biomass_g | Chr05K:44685227 | Chr05K | 44,685,227 | 0.4880 | < 0.001 | -0.7000 | 0.2167 | 0.0455 | 3.5260 | significant SNP | 0.0917 | PA | Pavir.5KG485300 |  |
| Pseudolabrys | Chr05K:44685227 | Chr05K | 44,685,227 | 0.4880 | < 0.001 | 0.8579 | 0.2368 | 0.0569 | 4.0592 | significant SNP | 0.0907 | PA | Pavir.5KG485300 |  |
| dry_biomass_g | Chr05K:8895158 | Chr05K | 8,895,158 | 0.1760 | < 0.001 | 1.0829 | 0.2750 | 0.0669 | 5.0143 | significant SNP | 0.1027 | PA | Pavir.5KG119200 | GO:0009892, GO:0043170, GO:0008152, GO:0009987 |
| Pseudolabrys | Chr05K:8895158 | Chr05K | 8,895,158 | 0.1760 | < 0.001 | -1.1695 | 0.3075 | 0.0625 | 4.4295 | significant SNP | 0.0788 | PA | Pavir.5KG119200 | GO:0009892, GO:0043170, GO:0008152, GO:0009987 |
| dry_biomass_g | Chr05K:8895168 | Chr05K | 8,895,168 | 0.2360 | < 0.001 | 0.9066 | 0.2623 | 0.0519 | 3.9698 | significant SNP | 0.0787 | PA | Pavir.5KG119200 | GO:0009892, GO:0043170, GO:0008152, GO:0009987 |
| Pseudolabrys | Chr05K:8895168 | Chr05K | 8,895,168 | 0.2360 | < 0.001 | -1.0548 | 0.2850 | 0.0593 | 4.2168 | significant SNP | 0.0701 | PA | Pavir.5KG119200 | GO:0009892, GO:0043170, GO:0008152, GO:0009987 |
| dry_biomass_g | Chr05N:52350110 | Chr05N | 52,350,110 | 0.3000 | < 0.001 | 0.8536 | 0.2562 | 0.0483 | 3.7203 | significant SNP | 0.0672 | PA | Pavir.5NG454900 |  |
| Pseudolabrys | Chr05N:52350110 | Chr05N | 52,350,110 | 0.3000 | < 0.001 | -1.0119 | 0.2987 | 0.0500 | 3.6056 | significant SNP | 0.0621 | PA | Pavir.5NG454900 |  |
| Actinobacteriota | Chr06K:6139838 | Chr06K | 6,139,838 | 0.2282 | < 0.001 | -0.5234 | 0.1529 | 0.0690 | 3.1978 | significant SNP | 0.0801 | NJ | Pavir.6KG071170 | GO:0032259, GO:0008152 |
| Pseudonocardia | Chr06K:6139838 | Chr06K | 6,139,838 | 0.2282 | < 0.001 | -0.7682 | 0.2031 | 0.0835 | 5.3769 | significant SNP | 0.0887 | NJ | Pavir.6KG071170 | GO:0032259, GO:0008152 |
| dry_biomass_g | Chr07K:32815708 | Chr07K | 32,815,708 | 0.0598 | < 0.001 | 1.1399 | 0.3455 | 0.0522 | 3.8156 | significant SNP | 0.0720 | NY | Pavir.7KG162000 | GO:0006464, GO:0044237, GO:0043170, GO:0008152 |
| Methylomirabilota | Chr07K:32815708 | Chr07K | 32,815,708 | 0.0598 | < 0.001 | 0.3370 | 0.0912 | 0.0650 | 4.3238 | significant SNP | 0.0639 | NY | Pavir.7KG162000 | GO:0006464, GO:0044237, GO:0043170, GO:0008152 |
| dry_biomass_g | Chr07K:32815708 | Chr07K | 32,815,708 | 0.0583 | < 0.001 | 1.7796 | 0.5192 | 0.0691 | 4.6496 | significant SNP | 0.0727 | NJ | Pavir.7KG162000 | GO:0006464, GO:0044237, GO:0043170, GO:0008152 |
| dry_biomass_g | Chr07K:48361582 | Chr07K | 48,361,582 | 0.2320 | < 0.001 | 0.7120 | 0.2204 | 0.0455 | 3.5259 | significant SNP | 0.0640 | PA | Pavir.7KG413400 | GO:0006928, GO:0044699, GO:0008152 |
| Pseudolabrys | Chr07K:48361582 | Chr07K | 48,361,582 | 0.2320 | < 0.001 | -0.8534 | 0.2480 | 0.0515 | 3.7060 | significant SNP | 0.0605 | PA | Pavir.7KG413400 | GO:0006928, GO:0044699, GO:0008152 |
| dry_biomass_g | Chr09N:66779803 | Chr09N | 66,779,803 | 0.2087 | < 0.001 | -1.1853 | 0.3380 | 0.0723 | 4.8439 | significant SNP | 0.0762 | NJ | Pavir.9NG675000 |  |
| Pseudonocardia | Chr09N:66779803 | Chr09N | 66,779,803 | 0.2087 | < 0.001 | -0.6411 | 0.2046 | 0.0581 | 3.8453 | significant SNP | 0.0606 | NJ | Pavir.9NG675000 |  |

**Table 3:** Model fit and test statistics for the generalized linear mixed models describing rhizo-sphere bacterial composition. Table includes Akaike information criterion, degrees of freedom for each model, and p-values for likelihood-ratio tests comparing Site-only and PC-only models to the Full models, respectively. (p >= 0.001 (***))

## Discussion

Against the background of major turnover in host-associated microbiomes across differing sites and difficulty in distinguishing between microbes that are truly host-associated or simply host-adjacent, it is often unclear to what extent host genetic variation impacts the composition of these diverse communities (Escudero-Martinez & Bulgarelli, 2019). Our study demonstrates the complexity of understanding the relationships between a uniquely large number of host plants sourced from across much of their natural range and their associated rhizosphere microbiomes at three divergent sites. A recent analysis of the switchgrass microbiome across its natural range showed that relative abundances of bacteria were strongly influenced by the interaction of host genetic variation and field site (Edwards et al., 2023). Similarly, our results suggest that switchgrass genetic variation can differentially influence microbiome composition between field trial sites and at different bacterial taxonomic scales. Our findings suggest that switchgrass rhizosphere microbiome composition may not only respond to host genetic influence at the ASV level but also at higher taxonomic levels in different sites, indicating that host genetic influence on microbiome composition may occur at both narrow and broad taxonomic scales. While not investigated here, this interaction between microbiome, site, and host genotypes could reflect (unknown) conserved microbial traits involved in response to genetic variation in host traits as well as differences in soil composition and other environmental factors among sites (Escudero-Martinez & Bulgarelli, 2019).

### Site effects on switchgrass microbiome composition

Each common garden in this study represents a unique soil type and management. The NY common garden presented a clay loam soil type and reduced management style. The NJ common garden was indicative of a well-suited (sandy, well-drained) and agriculturally managed (fertilizer was applied during establishment) soil type for switchgrass, while the PA common garden represented a poor (shale, rock, abandoned strip mine) and unmanaged soil type for switchgrass. These coarse abiotic differences ranged dramatically in both nutrient content and potential stressors, which likely influenced the differences in microbiome diversity and composition we observed between sites. Due to inflated zero values among rare taxa, which skew the distribution of microbial count data, bacterial diversity data rarely conform to the assumptions of MANOVA-like procedures. A non-parametric, one-way analysis of variance (NPMANOVA) was therefore used here and is preferred since it is tolerant of non-independent observations (e.g., microbe-microbe interactions) (Anderson, 2001). At the ASV level, we unsurprisingly found significant differences in Bray-Curtis dissimilarities of microbiome samples collected from each of the common gardens, illustrating distinct rhizosphere communities between them, likely with some important site-specific effects on microbiome assembly at the ASV level (Figure 2). Likewise, Shannon diversity of ASVs was differentiated between sites with the PA common garden exhibiting more diversity overall (Figure 1).

Vegetatively propagated plant material from the original study (Sutherland et al., 2022), may have had harbored remnant microbes within the root systems of the plants when they were transplanted to the three new sites used in this study. Therefore, we might presume to observe similar taxa in the switchgrass rhizosphere between sites, or even between studies. Technical limitations prevented us from directly comparing ASVs in the original study to ASVs in the common gardens presented here, but Marschner & Rumberger demonstrated that bacterial turnover can be relatively fast when introduced to new environments (2004). Thus, ASV composition between sites may differ after adapting to their new environments, and/or by the introduction of new ASVs into the host-associated rhizosphere, and, indeed, we observed that the site impact on ASV composition and diversity was significant.

To demonstrate this distinction, we observed similarity in the prevalence of certain taxa between common gardens when ASVs were labeled with the SILVA 138 taxonomic database and compared at higher taxonomic levels, even though compositional differences at the ASV-level between sites persisted (Figure 2). For example, the genus *Sphingomonas* was found in more than 90% of genotypes across all common gardens (Supplemental Figure 2), and yet the prevalence of many ASVs belonging to the *Sphingomonas* genus differed by common gardens (Supplemental Figure 4). This higher-level taxonomic similarity between sites could suggest a host genetic component to allowing specific microbial lineages to widely persist within the switchgrass rhizosphere where others do not. Given these differences in microbiome composition and diversity between growing sites, we then investigated how host genetic influence changes between growing sites by linking the relative abundance of microbial ASVs (and their respective higher taxonomic classifications) to switchgrass genetic variation.

### Host genetic effects on switchgrass microbiome composition

First, we employed broad-sense heritability (*H^2^*) to estimate the overall genetic effects of switchgrass on microbial relative abundances. Heritability is a concept in quantitative genetics that is used to describe the portion of a trait that can be explained by genetic variation. When applied to host-associated microbiomes, heritability can also be considered a metric of how community assembly is determined by host genetic variation (Sutherland et al., 2022; Wagner, 2021). Since *H^2^* estimates are sensitive to population structure, we estimated *H^2^* for each common garden individually (NJ, NY, PA) and then for host genotypes across sites (ALL). A significant reduction in the mean *H^2^* estimate for ALL was observed, as expected, likely due to the increased environmental variability. However, the mean estimate for ALL was ∼16%, indicating that host genetic influence persisted across sites.

One might hypothesize that stressed plants would devote more resources to microbial recruitment to account for stress – the ‘cry-for-help’ hypothesis (Rizaludin et al., 2021). While we were unable to explicitly test this hypothesis here, we believe this line of inquiry could be of interest to future studies using the switchgrass model. For instance, the NY common garden exhibited comparatively smaller *H^2^* estimates for bacterial abundances in the switchgrass rhizosphere (Figure 3). The NY site also had the highest plant biomass yield and most severe anthracnose symptoms on average, compared to PA or NJ (Supplemental Figure 3). Plants at the PA site were likely the most abiotically stressed due to site conditions (<u>See Materials and Methods</u>) and thus, as expected, exhibited the least biomass yield overall. *H^2^*estimates for bacterial abundances were generally the largest at the NJ site (Figure 3) where plants qualitatively suffered greatly from stem lodging. The NJ and PA sites showed comparable anthracnose symptoms, both of which were less severe than in NY (Supplemental Figure 3). Thus, the relative increase of heritability estimates for bacterial abundances in the switchgrass rhizosphere between sites appears to be more related to abiotic stress rather than to disease pressure. However, it is ultimately unclear whether biotic stress on switchgrass impacted host genetic influence on rhizosphere microbiome composition.

Next, we determined whether the heritability estimates were conserved within the bacterial phylogeny. This finding would suggest that the microbial traits that interact with genetic variation in switchgrass are phylogenetically conserved, and genetic variation in switchgrass is linked to selection for specific bacterial lineages. Phylogenetic conservatism is the tendency for traits of related species to resemble each other more than species drawn at random from the same tree. We hypothesized that host genetic influence might be conserved within regions of the microbial phylogeny where (unknown) functional traits are conserved. We observed a phylogenetic signal for heritability conserved at different regions of the bacterial phylogenetic tree. However, the pattern of conservation was unique to each of the common gardens, with NY having the most genera with a significant cluster (n =8) (Figure 4). Additionally, there was no overlap in which genera exhibited phylogenetic signals for heritability between sites. This suggests a host genetic influence on specific bacterial taxonomic groups due to their conserved traits, but that this influence may be contingent on the local species pool or environmental context. We want to highlight that we used relative microbial abundance data in this study (like most microbiome studies) rather than absolute microbial abundance data. It is important to note that the heritability of relative abundance is not necessarily the same as the heritability of absolute microbe abundance (Bruijning et al., 2023). However, the heritability and relative abundance of different taxa remain relevant to determining the relative advantages of certain microbes over others across host genetic differences (Bergelson et al., 2021; Deng et al., 2021; Wagner, 2021).

The interdependence of host genetics and growing site on microbiome assembly is widely hypothesized, and our results support this hypothesis in switchgrass. So, to explicitly test whether microbial assembly is driven by the shared response to host genotype and site, we implemented hierarchical distribution modeling. Our aim was two-fold. First, we wanted to test whether including site and SNP PCs (the full model) improved model fit compared to models that only included site or SNP PCs, respectively. Second, we tested whether grouping the random effects on ASV counts at different taxonomic levels improved model fit. By comparing model fits of the full, site-only, and PC-only models at different taxonomic levels, we could assess the strength of the genetic component influencing microbiome composition at different taxonomic levels.

Generally, the more complex models, with ASVs grouped at lower taxonomic levels, provided the best model fit. Regarding our first aim, we concluded that the full model was a better fit for the ASV counts data than site data or SNP PCs separately. We then compared the relative quality of each of the Full models with ASVs nested at different taxonomic levels. Model quality generally improved when ASV counts were grouped at lower taxonomic levels. The full unnested model was relatively the best fit for the data (Table 3). These results suggest that the relationship between host genotype and site at the ASV level best explains variance in switchgrass rhizosphere microbiome composition. However, the addition of host genetic information to site data significantly improved model fit at all taxonomic levels, suggesting that genome-wide host genetic influence on microbial taxa also exists. Therefore, our results suggest that host genetic influence on microbiome composition exists at all taxonomic levels, but is stronger at lower levels (genus; ASV).

### Genotype-by-Environment-by-Microbiome associations with switchgrass performance traits

We found that the relationships between the relative abundance of genera in the switchgrass rhizosphere and biomass yield and anthracnose disease severity were site-specific. Notably, in the NJ common garden, where nitrogen fertilizer was applied during plant establishment, we observed enrichment for the genus *Nitrospira* compared to the other sites. *Nitrospira* is an aerobic chemolithoautotrophic genus, containing a high proportion of bacteria that are known to oxidize ammonia in soils and release nitrogen to the atmosphere as nitrous-oxide (N_2_O) and/or dinitrogen (N_2_) (Daims & Wagner, 2018). We observed a negative relationship (r = -0.357, p_adj_ < 0.001) between the relative abundance of *Nitrospira* and plant biomass at the NJ common garden, but not at the other sites. One possibility is a competitive dynamic between switchgrass hosts and *Nitrospira* in the presence of supplementary nitrogen (Kuzyakov & Xu, 2013), which could impede biomass yield.

At the PA common garden, we observed a negative relationship between plant biomass yield and the relative abundance of the MND1 genus (r = -0.299, padj < 0.001). MND1 belongs to the family Nitrosomonadaceae and, much like *Nitrospira*, is known to contain ammonia-oxidizing bacteria (Kong, Wang, Niu, & Miao, 2016). Moreover, there was a positive relationship between the relative abundance of MND1 and anthracnose disease severity (r = 0.284, padj < 0.001). This inverse relationship between biomass production and anthracnose disease severity for plants enriched for MND1 suggests that MND1 may play a role in the pathogenicity of anthracnose or that MND1 responds to host anthracnose symptoms. Alternatively, much like at the NJ site, competition for available nitrogen could potentially have an impact on switchgrass yield. Future work examining isotope-labeled nitrogen assimilation in switchgrass, controlling for microbiome composition and infection, could confirm these hypotheses.

Finally, we conducted a genome-wide association study (GWAS) to identify shared loci associated with the relative abundance of bacterial taxa, biomass yield, and anthracnose disease severity at the common gardens. We observed overlapping loci associated with these traits and the abundance of seven bacterial taxa (Table 2). Most of the SNPs were in genes (n = 6) associated with gene ontologies related to plant metabolism: GO:0008152. While the pathways for these genes require further investigation, we can gain some insights from these relationships that could positively impact host genetic control over certain microbiota in the switchgrass rhizosphere. For instance, *Pseudolabrys* is a genus of bacteria from the Nitrobacteraceae family. At PA, *Pseudolabrys* is negatively correlated with biomass yield and SNP PC1, and positively correlated with SNP PC2 (Table 1). SNP PC1 and PC2 capture the coarse allelic variation associated with switchgrass ecotype (Supplemental Figure 1). In addition, SNPs with effects that positively correlate with biomass yield at PA, negatively correlate with *Pseudolabrys*, and vice-versa (Table 2, Supplemental Figure 5). Therefore, these genes (Pavir.1KG354700, Pavir.3KG080100, Pavir.4KG003200, Pavir.4NG164500, Pavir.5KG485300, Pavir.5KG119200, Pavir.5KG119200, Pavir.5NG454900, Pavir.7KG413400) could offer potential pathways for improving switchgrass biomass yield grown on marginal lands by influencing the abundance of certain bacteria in the rhizosphere associated with this trait.

## Conclusion

We observed, as expected, a strong site influence over the switchgrass bacterial rhizosphere microbiome, likely driven by differences in the abiotic features in each common garden and their respective bacterial species pools during plant establishment. Additionally, host genetic influence differed between sites and between phylogenetic clades for bacteria in the rhizosphere. Taken together, the relationships between host genotype, environmental features, and the plant-associated microbiome are highly contextual, depending on several ecological and evolutionary factors. Future work might continue to consider the interdependence of these factors to better understand the influences shaping rhizosphere microbiome composition in switchgrass.

## Supplemental Figures

**Supplemental Figure 1: Switchgrass SNP PCA Describes Ecotype and Geographic Divergence**

Principal component analysis of switchgrass SNPs characterizes genetic divergence between Lowland (gold), Intermediate (light blue), and Upland (black) switchgrass ecotypes and geographic regions. Lowland ecotype divergence along PC2 is characterized by the population divergence between the South population and Northeast Population, represented here by ellipses following a multivariate t-distribution from the centroid for the North (orange), Northeast (dark red), South (green), West (olive), and East (pink) populations.

**Supplemental Figure 2: Bacterial Prevalence by Taxonomic Level**

Bacterial prevalence for all genotypes between common gardens represented for each taxonomic level. Taxa with prevalence less than 25% removed for clarity. Dashed-line at 90% prevalence.

**Supplemental Figure 3: Mean Biomass and Anthracnose Ratings by Site**

The mean biomass measurements in grams (A) and mean anthracnose disease ratings (B) for NJ (red), NY, (green), and PA (yellow).

**Supplemental Figure 4: Sphingomonas ASV Prevalence by Site**

The prevalence of ASVs assigned to the Sphingomonas genus greater than 5% among genotypes at ALL sites (A), NY (B), PA (C), and NJ (D) common gardens. Dashed-line at 90% prevalence.

**Supplemental Figure 5: Shared SNPs and Effect Sizes for Biomass Yield and Pseudolabrys relative abundances at the PA common garden**

Effect sizes for SNPs associated with both Biomass yield (g, green) and Pseudolabrys relative abundances (red) at the PA common garden.

## References

1. Ackerly, D. D. (2003). Community assembly, niche conservatism, and adaptive evolution in changing environments. International Journal of Plant Sciences, 164(SUPPL. 3), 165–184. doi: 10.1086/368401/ASSET/IMAGES/LARGE/FG7.JPEG

2. Anderson, M. J. (2001). A new method for non-parametric multivariate analysis of variance. Austral Ecology, 26(1), 32–46. doi: 10.1111/j.1442-9993.2001.01070.pp.x

3. Apprill, A., Mcnally, S., Parsons, R., & Weber, L. (2015). Minor revision to V4 region SSU rRNA 806R gene primer greatly increases detection of SAR11 bacterioplankton. Aquatic Microbial Ecology. doi: 10.3354/ame01753

4. Bergelson, J., Brachi, B., Roux, F., & Vailleau, F. (2021). Assessing the potential to harness the microbiome through plant genetics. Current Opinion in Biotechnology, 70, 167–173. doi: 10.1016/j.copbio.2021.05.007

5. Bruijning, M., Ayroles, J. F., Henry, L. P., Koskella, B., Meyer, K. M., & Metcalf, C. J. E. (2023). Relative abundance data can misrepresent heritability of the microbiome. Microbiome, 11(1), 222. doi: 10.1186/s40168-023-01669-w

6. Brunel, C., Pouteau, R., Dawson, W., Pester, M., Ramirez, K. S., & van Kleunen, M. (2020). Towards Unraveling Macroecological Patterns in Rhizosphere Microbiomes. Trends in Plant Science, 25(10), 1017–1029. doi: 10.1016/J.TPLANTS.2020.04.015

7. Brzyski, D., Peterson, C. B., Sobczyk, P., Candès, E. J., Bogdan, M., & Sabatti, C. (2017). Controlling the rate of GWAS false discoveries. Genetics. doi: 10.1534/genetics.116.193987

8a. Callahan, B. J., McMurdie, P. J., Rosen, M. J., Han, A. W., Johnson, A. J. A., & Holmes, S. P. (2016). DADA2: High-resolution sample inference from Illumina amplicon data. Nature Methods, 13(7), 581–583. doi: 10.1038/nmeth.3869

8. Compant, S., Clément, C., & Sessitsch, A. (2010, May 1). Plant growth-promoting bacteria in the rhizo-and endosphere of plants: Their role, colonization, mechanisms involved and prospects for utilization. Soil Biology and Biochemistry, Vol. 42, pp. 669–678. doi: 10.1016/j.soilbio.2009.11.024

9. Csikós, N., & Tóth, G. (2023). Concepts of agricultural marginal lands and their utilisation: A review. Agricultural Systems, 204, 103560. doi: 10.1016/J.AGSY.2022.103560

10. Daims, H., & Wagner, M. (2018). Nitrospira. Trends in Microbiology, 26(5), 462–463. doi: 10.1016/J.TIM.2018.02.001

11. Danecek, P., Bonfield, J. K., Liddle, J., Marshall, J., Ohan, V., Pollard, M. O., … Li, H. (2021). Twelve years of SAMtools and BCFtools. GigaScience, 10(2), 1–4. doi: 10.1093/GIGASCIENCE/GIAB008

12. Delgado-Baquerizo, M., Reich, P. B., Khachane, A. N., Campbell, C. D., Thomas, N., Freitag, T. E., … Singh, B. K. (2017). It is elemental: soil nutrient stoichiometry drives bacterial diversity. Environmental Microbiology, 19(3), 1176–1188. doi: 10.1111/1462-2920.13642

13. Deng, S., Caddell, D. F., Xu, G., Dahlen, L., Washington, L., Yang, J., & Coleman-Derr, D. (2021). Genome wide association study reveals plant loci controlling heritability of the rhizosphere microbiome. ISME Journal, 15(11). doi: 10.1038/s41396-021-00993-z

14. Edwards, J. A., Bishnoi Saran, U., Bonnette, J., MacQueen, A., Yin, J., uyen Nguyen, T., … Juenger, T. E. (2023). Genetic determinants of switchgrass-root-associated microbiota in field sites spanning its natural range. doi: 10.1016/j.cub.2023.03.078

15. Escudero-Martinez, C., & Bulgarelli, D. (2019). Tracing the evolutionary routes of plant– microbiota interactions. Current Opinion in Microbiology, 49, 34–40. doi: 10.1016/J.MIB.2019.09.013

16. Evans, J., Sanciangco, M. D., Lau, K. H., Crisovan, E., Barry, K., Daum, C., … Buell, C. R. (2018). Extensive Genetic Diversity is Present within North American Switchgrass Germplasm. The Plant Genome, 11(1), 1–16. doi: 10.3835/plantgenome2017.06.0055

17. Fierer, N., Ladau, J., Clemente, J. C., Leff, J. W., Owens, S. M., Pollard, K. S., … McCulley, R. L. (2013). Reconstructing the microbial diversity and function of pre-agricultural tallgrass prairie soils in the United States. Science, 342(6158), 621–624. doi: 10.1126/SCIENCE.1243768/SUPPL_FILE/FIERER.SM.PDF

18. Hartman, J. C., Nippert, J. B., Orozco, R. A., & Springer, C. J. (2011). Potential ecological impacts of switchgrass (Panicum virgatum L.) biofuel cultivation in the Central Great Plains, USA. Biomass and Bioenergy, 35(8), 3415–3421. doi: 10.1016/J.BIOMBIOE.2011.04.055

19. Hestrin, R., Lee, M. R., Whitaker, B. K., & Pett-Ridge, J. (2021). The switchgrass microbiome: A review of structure, function, and taxonomic distribution. Phytobiomes Journal. doi: 10.1094/pbiomes-04-20-0029-fi

20. Hothorn, A. Z. and T. (2002). Diagnostic Checking in Regression Relationships. R News, 2(3), 7– 10.

21. Hu, L., Robert, C. A. M., Cadot, S., Zhang, X., Ye, M., Li, B., … Erb, M. (2018). Root exudate metabolites drive plant-soil feedbacks on growth and defense by shaping the rhizosphere microbiota. Nature Communications, 9(1), 1–13. doi: 10.1038/s41467-018-05122-7

22. Ihaka, R., & Gentleman, R. (1996). R: A Language for Data Analysis and Graphics. Journal of Computational and Graphical Statistics, 5(3), 299–314. doi: 10.1080/10618600.1996.10474713

23. Kang, H. M., Sul, J. H., Service, S. K., Zaitlen, N. A., Kong, S. Y., Freimer, N. B., … Eskin, E. (2010). Variance component model to account for sample structure in genome-wide association studies. Nature Genetics, 42(4), 348–354. doi: 10.1038/ng.548

24. Keck, F., Rimet, F., Bouchez, A., & Franc, A. (2016). phylosignal: an R package to measure, test, and explore the phylogenetic signal. Ecology and Evolution, 6(9), 2774–2780. doi: 10.1002/ECE3.2051

25. Kembel, S. W., & Hubbell, S. P. (2006). THE PHYLOGENETIC STRUCTURE OF A NEOTROPICAL FOREST TREE COMMUNITY. Ecology, 87(7), 86–99. doi: 10.1890/0012-9658

26. Kong, Q., Wang, Z. bin, Niu, P. fei, & Miao, M. sheng. (2016). Greenhouse gas emission and microbial community dynamics during simultaneous nitrification and denitrification process. Bioresource Technology, 210, 94–100. doi: 10.1016/J.BIORTECH.2016.02.051

27. Kuzyakov, Y., & Xu, X. (2013). Competition between roots and microorganisms for nitrogen: mechanisms and ecological relevance. New Phytologist, 198(3), 656–669. doi: 10.1111/NPH.12235

28. Lasky, J. R., Keitt, T. H., Weeks, B. C., & Economo, E. P. (2017). A hierarchical model of whole assemblage island biogeography. Ecography, 40(8), 982–990. doi: 10.1111/ECOG.02303

29. Li, H. (2013). Aligning sequence reads, clone sequences and assembly contigs with BWA-MEM. Retrieved from https://arxiv.org/abs/1303.3997v2

30. Lowry, D. B., Behrman, K. D., Grabowski, P., Morris, G. P., Kiniry, J. R., & Juenger, T. E. (2015). Adaptations between Ecotypes and along Environmental Gradients in Panicum virgatum*. *183*(5), 682–692. doi: 10.1086/675760

31. Lu, F., Lipka, A. E., Glaubitz, J., Elshire, R., Cherney, J. H., Casler, M. D., … Costich, D. E. (2013). Switchgrass Genomic Diversity, Ploidy, and Evolution: Novel Insights from a Network-Based SNP Discovery Protocol. PLoS Genetics. doi: 10.1371/journal.pgen.1003215

32a. Mahmud, K., Missaoui, A., Lee, K., Ghimire, B., Presley, H. W., & Makaju, S. (2021). Rhizosphere microbiome manipulation for sustainable crop production. Current Plant Biology, 27, 100210. doi: 10.1016/J.CPB.2021.100210

32. Mansfield, J., Genin, S., Magori, S., Citovsky, V., Sriariyanum, M., Ronald, P., … Foster, G. D. (2012). Top 10 plant pathogenic bacteria in molecular plant pathology. Molecular Plant Pathology, 13(6), 614–629. doi: 10.1111/J.1364-3703.2012.00804.X

33. Marschner, P., & Rumberger, A. (2004). Rapid changes in the rhizosphere bacterial community structure during re-colonization of sterilized soil. Biology and Fertility of Soils, 40(1), 1–6. doi: 10.1007/S00374-004-0736-4/FIGURES/3

34. Martin, M. (2011). Cutadapt removes adapter sequences from high-throughput sequencing reads. *EMBnet.Journal*, 17(1), 10–12. doi: 10.14806/EJ.17.1.200

35. Martiny, J. B. H. H., Jones, S. E., Lennon, J. T., & Martiny, A. C. Microbiomes in light of traits: A phylogenetic perspective. , 350 Science § (2015).

36. McMurdie, P. J., & Holmes, S. (2013). phyloseq: An R Package for Reproducible Interactive Analysis and Graphics of Microbiome Census Data. PLoS ONE, 8(4), e61217. doi: 10.1371/journal.pone.0061217

37. Oksanen, J., Blanchet, F. G., Friendly, M., Kindt, R., Legendre, P., Mcglinn, D., … Maintainer, H. W. (2019). vegan: Community Ecology Package. R package version 2.5-5. https://CRAN.R-project.org/package=vegan. *Community Ecology Package*.

38. Parada, A. E., Needham, D. M., & Fuhrman, J. A. (2016). Every base matters: Assessing small subunit rRNA primers for marine microbiomes with mock communities, time series and global field samples. Environmental Microbiology. doi: 10.1111/1462-2920.13023

39. Peiffer, J. A., Spor, A., Koren, O., Jin, Z., Tringe, S. G., Dangl, J. L., … Ley, R. E. (2013). Diversity and heritability of the maize rhizosphere microbiome under field conditions. Proceedings of the National Academy of Sciences of the United States of America, 110(16), 6548–6553. doi: 10.1073/PNAS.1302837110/SUPPL_FILE/PNAS.201302837SI.PDF

40. Pérez-Jaramillo, J. E., Mendes, R., & Raaijmakers, J. M. (2016). Impact of plant domestication on rhizosphere microbiome assembly and functions. Plant Molecular Biology, 90(6), 635–644. doi: 10.1007/s11103-015-0337-7

41. Prosser, J. I., & Martiny, J. B. H. (2020). Conceptual challenges in microbial community ecology. Philosophical Transactions of the Royal Society B, 375(1798). doi: 10.1098/RSTB.2019.0241

42. Quast, C., Pruesse, E., Yilmaz, P., Gerken, J., Schweer, T., Yarza, P., … Glö Ckner, F. O. (n.d.). The SILVA ribosomal RNA gene database project: improved data processing and web-based tools. doi: 10.1093/nar/gks1219

43. Rizaludin, M. S., Stopnisek, N., Raaijmakers, J. M., & Garbeva, P. (2021). The Chemistry of Stress: Understanding the ‘Cry for Help’ of Plant Roots. Metabolites 2021, *Vol. 11, Page 357*, *11*(6), 357. doi: 10.3390/METABO11060357

44. Rossum, B. van, Kruijer, W., Eeuwijk, F. van, & Boer, M. (2020). Package “statgenGWAS” Title Genome Wide Association Studies. doi: 10.1038/ng.548

45. Ruhl, I. A., Sheremet, A., Smirnova, A. V., Sharp, C. E., Grasby, S. E., Strous, M., & Dunfield, P. F. (2022). Microbial Functional Diversity Correlates with Species Diversity along a Temperature Gradient. MSystems, 7(1). doi: 10.1128/MSYSTEMS.00991-21/SUPPL_FILE/MSYSTEMS.00991-21-ST004.PDF

46. Schloss, P. D., & Arbor, A. (2023). Rarefaction is currently the best approach to control for uneven sequencing effort in amplicon sequence analyses. *BioRxiv*, 2023.06.23.546313.doi: 10.1101/2023.06.23.546313

47. Serna-Chavez, H. M., Fierer, N., & Van Bodegom, P. M. (2013). Global drivers and patterns of microbial abundance in soil. Global Ecology and Biogeography, 22(10), 1162–1172. doi: 10.1111/GEB.12070

48. Simon Andrews. (2020). Babraham Bioinformatics - FastQC A Quality Control tool for High Throughput Sequence Data. *Soil*, Vol. 5, pp. 47–81. Retrieved from https://www.bioinformatics.babraham.ac.uk/projects/fastqc/

49. Stewart, C. E., Roosendaal, D., Denef, K., Pruessner, E., Comas, L. H., Sarath, G., … Soundararajan, M. (2017). Seasonal switchgrass ecotype contributions to soil organic carbon, deep soil microbial community composition and rhizodeposit uptake during an extreme drought. Soil Biology and Biochemistry, 112, 191–203. doi: 10.1016/j.soilbio.2017.04.021

50. Sutherland, J., Bell, T., Trexler, R. V., Carlson, J. E., & Lasky, J. R. (2022). Host genomic influence on bacterial composition in the switchgrass rhizosphere. Molecular Ecology, 00, 1–17. doi: 10.1111/MEC.16549

51. Trexler, R. V., & Bell, T. H. (2019). Testing sustained soil-to-soil contact as an approach for limiting the abiotic influence of source soils during experimental microbiome transfer. FEMS Microbiology Letters. doi: 10.1093/femsle/fnz228

52. Ulbrich, T. C., Friesen, M. L., Roley, S. S., Tiemann, L. K., & Evans, S. E. (2021). Intraspecific variability in root traits and edaphic conditions influence soil microbiomes across 12 switchgrass cultivars. Phytobiomes Journal, 5(1), 108–120. doi: 10.1094/PBIOMES-12-19-0069-FI/ASSET/IMAGES/LARGE/PBIOMES-12-19-0069-FIT4.JPEG

53. VanRaden, P. M. (2008). Efficient methods to compute genomic predictions. Journal of Dairy Science, 91(11), 4414–4423. doi: 10.3168/jds.2007-0980

54. Wagner, M. R. (2021). Prioritizing host phenotype to understand microbiome heritability in plants. New Phytologist. doi: 10.1111/NPH.17622

55. Weiss, S., Xu, Z. Z., Peddada, S., Amir, A., Bittinger, K., Gonzalez, A., … Knight, R. (2017). Normalization and microbial differential abundance strategies depend upon data characteristics. Microbiome, 5(1), 27. doi: 10.1186/s40168-017-0237-y

56. Wright, E. S. (n.d.). Using DECIPHER v2.0 to Analyze Big Biological Sequence Data in R.

56a. Zhalnina, K., Louie, K. B., Hao, Z., Mansoori, N., Da Rocha, U. N., Shi, S., … Brodie, E. L. (2018). Dynamic root exudate chemistry and microbial substrate preferences drive patterns in rhizosphere microbial community assembly. Nature Microbiology. doi: 10.1038/s41564-018-0129-3

57. Zhu, Q., Mai, U., Pfeiffer, W., Janssen, S., Asnicar, F., Sanders, J. G., … Knight, R. (2019). Phylogenomics of 10,575 genomes reveals evolutionary proximity between domains Bacteria and Archaea. Nature Communications 2019 10:1, *10*(1), 1–14. doi: 10.1038/s41467-019-13443-4

