## Supplementary figures and images for "Bacterial assembly in the switchgrass rhizosphere is shaped by phylogeny, host genotype, and growing site"

### Supplemental Figure 1

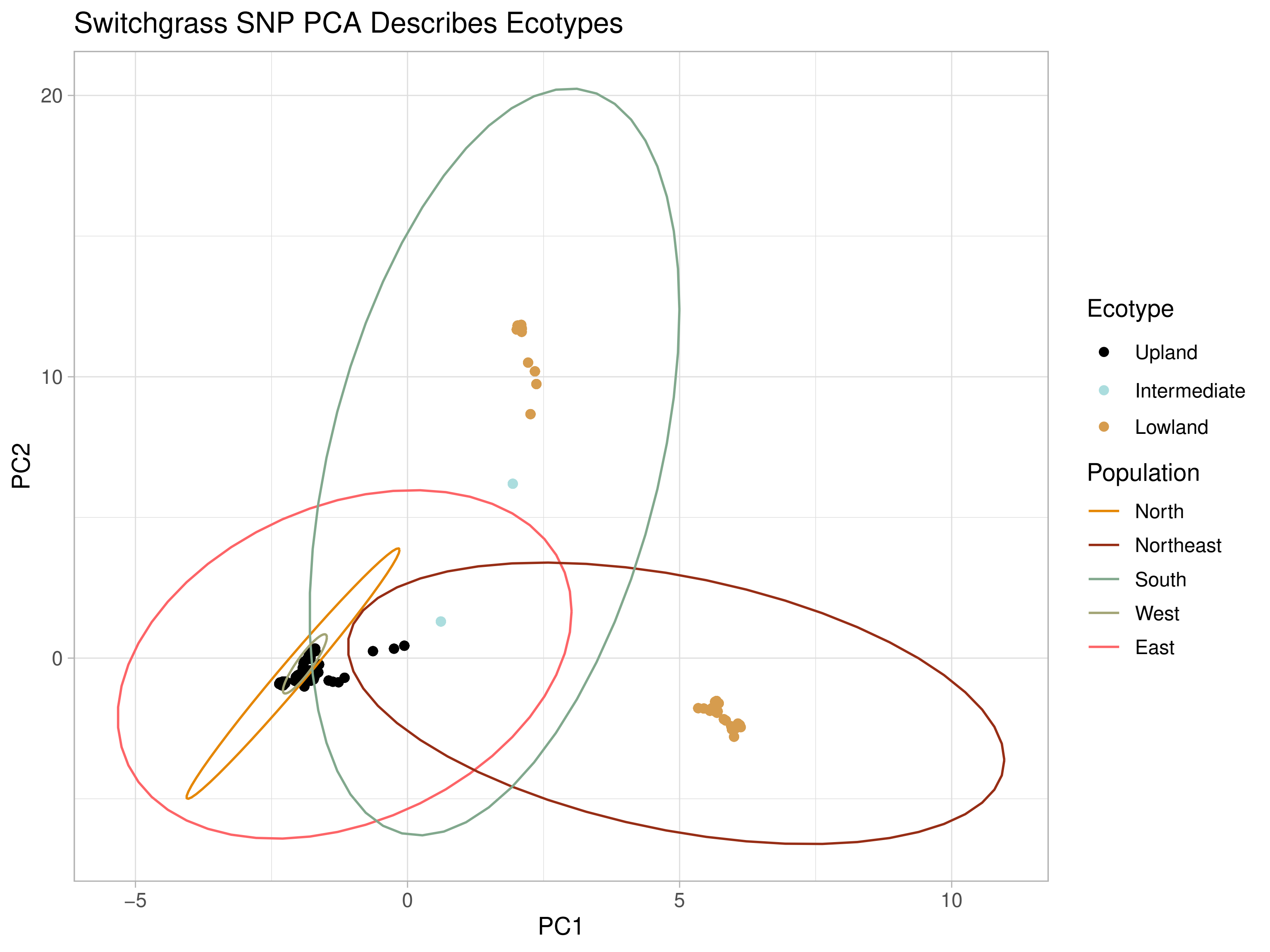

### Supplemental Figure 2

#### Bacterial Prevalence by Taxonomic Level

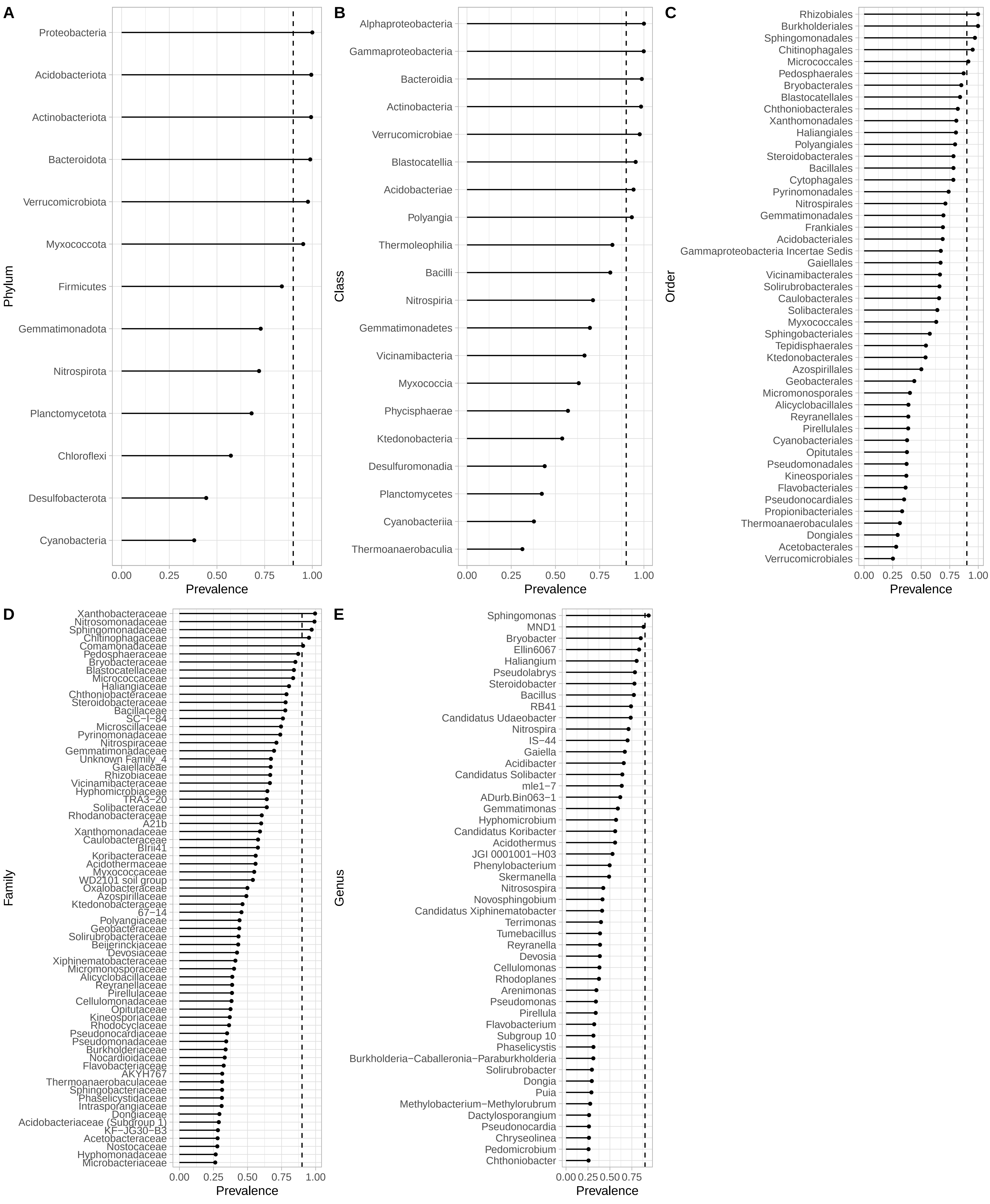

### Supplemental Figure 3

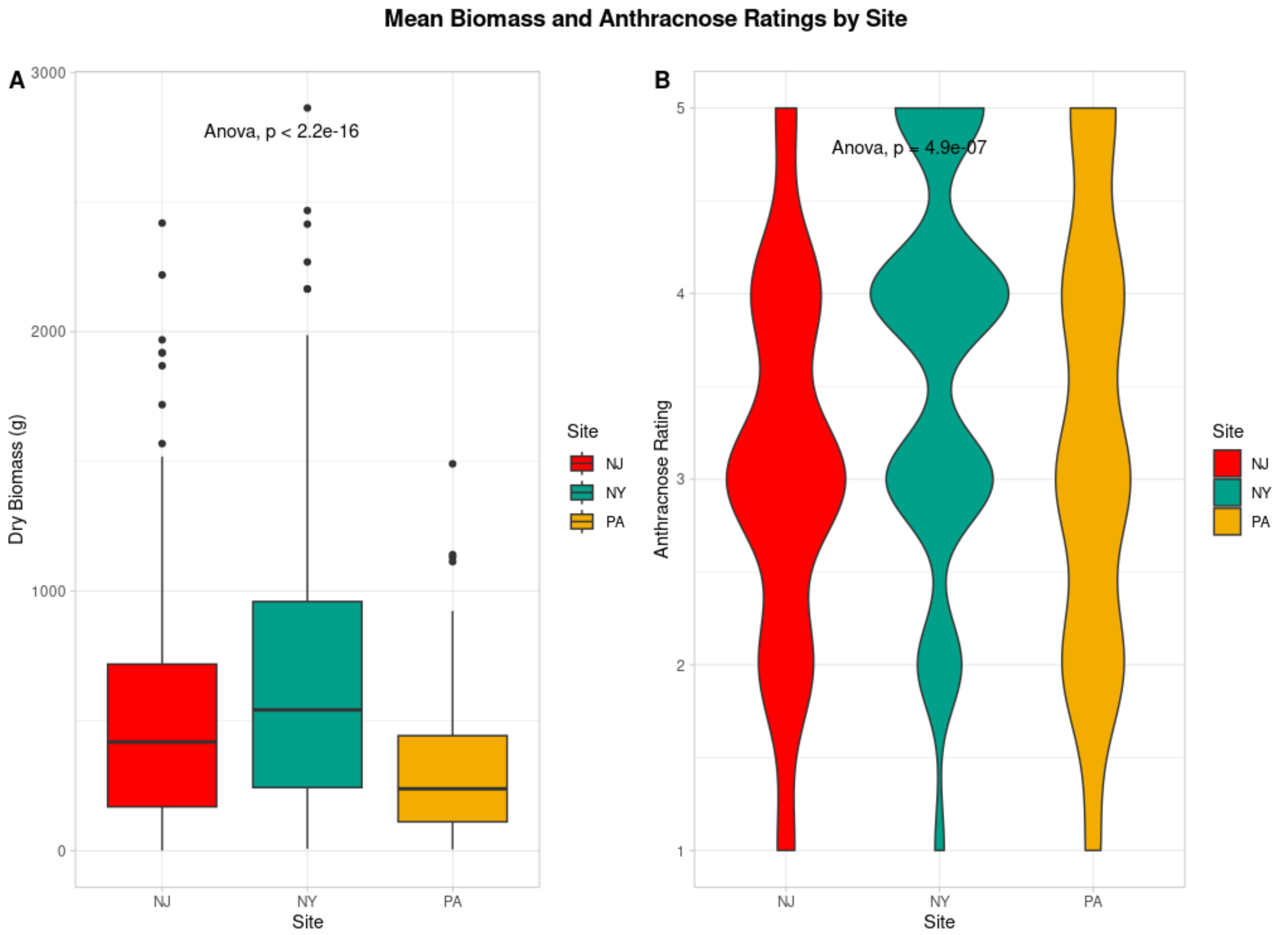

### Supplemental Figure 4

#### Sphingomonas Prevalence by Site

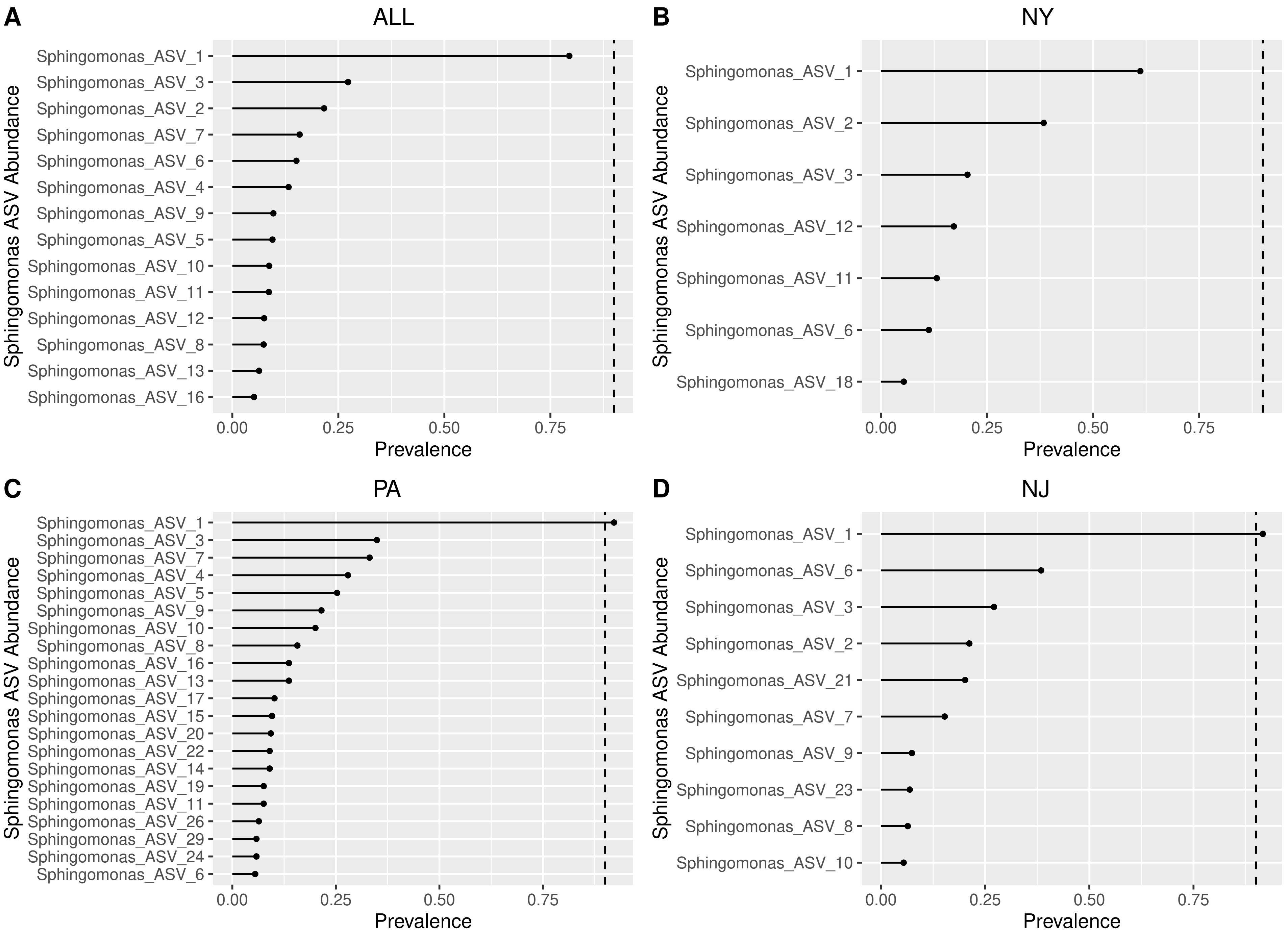

### Supplemental Figure 5

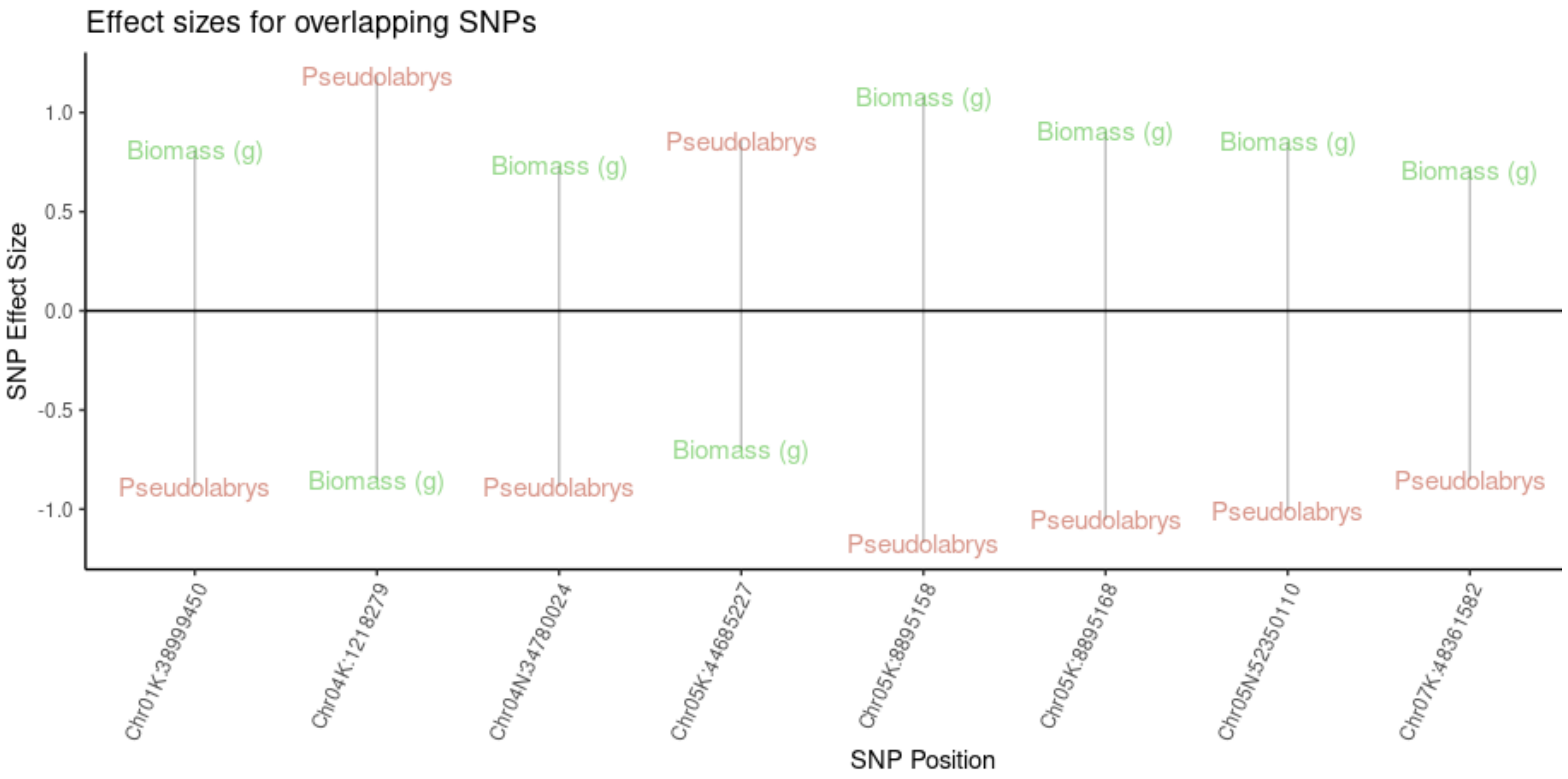
